# An integrative view of the regulatory and transcriptional landscapes in mouse hematopoiesis

**DOI:** 10.1101/731729

**Authors:** Guanjue Xiang, Cheryl A. Keller, Elisabeth Heuston, Belinda M. Giardine, Lin An, Alexander Q. Wixom, Amber Miller, April Cockburn, Michael E.G. Sauria, Kathryn Weaver, Jens Lichtenberg, Berthold Göttgens, Qunhua Li, David Bodine, Shaun Mahony, James Taylor, Gerd A. Blobel, Mitchell J. Weiss, Yong Cheng, Feng Yue, Jim Hughes, Douglas R. Higgs, Yu Zhang, Ross C. Hardison

## Abstract

Thousands of epigenomic datasets have been generated in the past decade, but it is difficult for researchers to effectively utilize all the data relevant to their projects. Systematic integrative analysis can help meet this need, and the VISION project was established for **V**al**I**dated **S**ystematic **I**ntegrati**ON** of epigenomic data in hematopoiesis. Here, we systematically integrated extensive data recording epigenetic features and transcriptomes from many sources, including individual laboratories and consortia, to produce a comprehensive view of the regulatory landscape of differentiating hematopoietic cell types in mouse. By employing IDEAS as our **I**ntegrative and **D**iscriminative **E**pigenome **A**nnotation **S**ystem, we identified and assigned epigenetic states simultaneously along chromosomes and across cell types, precisely and comprehensively. Combining nuclease accessibility and epigenetic states produced a set of over 200,000 candidate *cis*-regulatory elements (cCREs) that efficiently capture enhancers and promoters. The transitions in epigenetic states of these cCREs across cell types provided insights into mechanisms of regulation, including decreases in numbers of active cCREs during differentiation of most lineages, transitions from poised to active or inactive states, and shifts in nuclease accessibility of CTCF-bound elements. Regression modeling of epigenetic states at cCREs and gene expression produced a versatile resource to improve selection of cCREs potentially regulating target genes. These resources are available from our VISION website (usevision.org) to aid research in genomics and hematopoiesis.

## Introduction

Individual laboratories and major consortia (e.g., The ENCODE Project Consortium 2012; Cheng *et al*. 2014; Yue *et al*. 2014; Roadmap Epigenomics *et al*. 2015; Stunnenberg *et al*. 2016; The ENCODE Project Consortium *et al*. 2019) have produced thousands of genome-wide datasets on transcriptomes and many epigenetic features, including nuclease accessibility, histone modifications, and transcription factor occupancy, across diverse cell types. However, it is challenging for individual investigators to find all the data relevant to their projects or to incorporate the data effectively into analyses and hypothesis generation. One approach to address this challenge of overwhelming data is to integrate the deep and diverse datasets (Ernst and Kellis 2010; Ernst and Kellis 2012; Hoffman et al. 2012; Hoffman et al. 2013; Zhou and Troyanskaya 2015; Greenside et al. 2018; Lee et al. 2018; Ludwig et al. 2019). An effective integration will produce simplified representations of the data that facilitate discoveries and lead to testable hypotheses about functions of genomic elements and mechanisms of regulatory processes. Our multi-lab project called VISION (for **V**al**I**dated **S**ystematic **I**ntegrati**ON** of hematopoietic epigenomes) is endeavoring to meet this challenge by focusing on an important biological system, hematopoietic differentiation. Not only is hematopoietic differentiation an important biological and medical system with abundant epigenetic data available (e.g., Cheng *et al*. 2009; Fujiwara *et al*. 2009; Yu *et al*. 2009; Wilson *et al*. 2010; Pilon *et al*. 2011; Tijssen *et al*. 2011; Wong *et al*. 2011; Wu *et al*. 2011; Kowalczyk *et al*. 2012; Su *et al*. 2013; Lara-Astiaso *et al*. 2014; Pimkin *et al*. 2014; Wu *et al*. 2014; Corces *et al*. 2016; Huang *et al*. 2016; Heuston *et al*. 2018; Ludwig *et al*. 2019), but it also provides a powerful framework for validation of the integrative modeling. Specifically, work over prior decades has established key concepts that a successful modeling effort should recapitulate, and predictions of the modeling can be tested genetically in animals and cell lines. Here, we report on our initial systematic integrative modeling of mouse hematopoiesis.

The production of many distinct blood cell types from a common stem cell (hematopoiesis) is critically important for human health (Orkin and Zon 2008), and it has been studied intensively in humans and mouse. Despite some differences between these species (An et al. 2014; Cheng et al. 2014; Pishesha et al. 2014), the mouse system has served as a good model for many aspects of hematopoiesis in humans and mammals (Sykes and Scadden 2013). In adult mammals, all blood cells are produced from mesodermally-derived, self-renewing hematopoietic stem cells (HSCs) located in the bone marrow (Till and McCulloch 1961; Kondo et al. 2003). Studies of populations of multilineage progenitor cells, purified using cell surface markers (Weissman and Shizuru 2008), show that hematopoietic differentiation proceeds from HSC through progenitor cells with progressively more restricted lineage potential, eventually committing to a single cell lineage (Reya et al. 2001). More recent analyses of single cell transcriptomes have revealed heterogeneity in each of these cell populations (Sanjuan-Pla et al. 2013; Psaila et al. 2016). Overall, analyses of single cell transcriptomes support an ensemble of pathways for differentiation (Nestorowa et al. 2016; Laurenti and Gottgens 2018). Regardless of the complexity in cell-fate pathways, it is clear that changes in patterns of gene expression drive the differentiation program (Cantor and Orkin 2002; Graf and Enver 2009). Mis-regulation of gene expression patterns can cause diseases such as leukemias and anemias (Higgs 2013; Lee and Young 2013; Ling and Crispino 2020), and thus, efforts to better understand the molecular mechanisms regulating gene expression can help uncover the processes underlying cancers and blood disorders.

Comprehensive epigenomic and transcriptomic data can be used to describe how both the patterns of gene expression and the regulatory landscapes change during hematopoietic differentiation. Previous reports provided many insights and datasets on epigenomic changes during hematopoiesis in mouse (e.g., Lara-Astiaso et al. 2014) and in human (e.g., Adams et al. 2012; Corces et al. 2016). Additional informative datasets have come from detailed studies in cell line models of hematopoietic differentiation. In the intensively studied process of hematopoiesis, such comprehensive datasets could encompass virtually all the regulatory and transcriptional changes that occur during differentiation. However, distilling the regulatory events that are most critical to producing the transcriptional patterns needed for distinctive cell types is still a major challenge. Here, our major aim is to systematically integrate the extensive epigenomic data to improve accessibility and understanding of the data and to facilitate the generation of novel hypotheses about changes in the regulatory landscape during hematopoietic differentiation. We determined epigenetic states, which are common combinations of epigenetic features, to generate a readily interpretable “painting” of the epigenomic landscape across selected mouse hematopoietic cell populations. The state assignments coupled with peaks of nuclease accessibility produced an initial compendium of over 200,000 candidate *cis*-regulatory elements (cCREs) active in one or more hematopoietic lineages in mouse, which are valuable for further studies of hematopoietic gene regulation.

## Results

### Epigenomic and transcriptomic datasets of mouse hematopoietic cells

We reasoned that integrative analysis of the large number of genome-wide determinations of RNA levels and epigenetic features should provide an accessible view of the information that would help investigators utilize these diverse datasets, and it may lead to novel insights into gene regulation. To conduct the integrative and discriminative analysis, we collated the raw sequence data for 150 determinations of relevant epigenetic features (104 experiments after merging replicates) across 20 cell types or populations (Fig. 1A), including histone modifications and CTCF by ChIP-seq, nuclease accessibility of DNA in chromatin by ATAC-seq and DNase-seq, and transcriptomes by RNA-seq. The purified cell populations and cell lines are described in detail in the Supplemental Material, section1.

**Figure 1.**
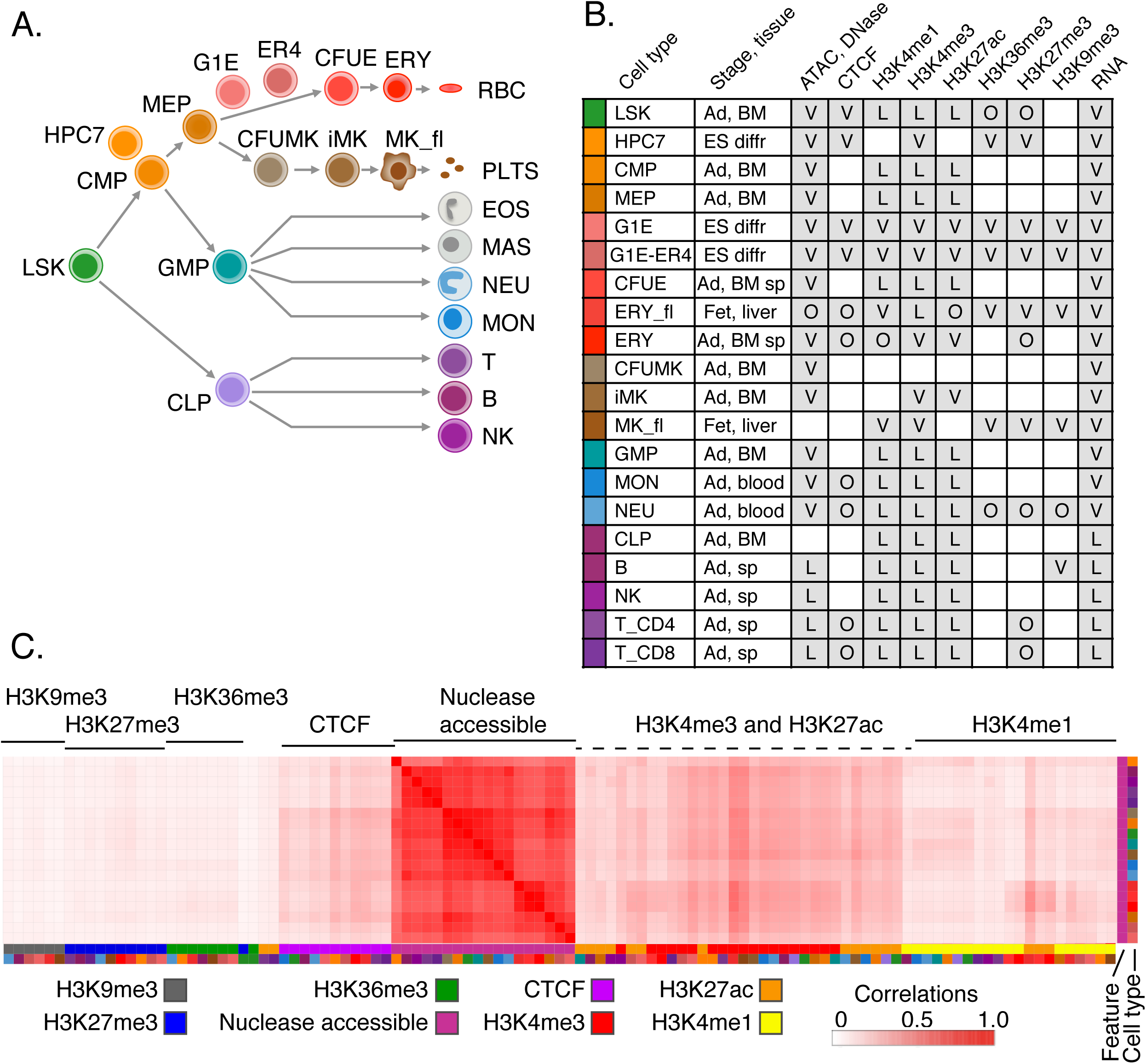
Hematopoietic cell types and datasets used for integrative analysis. **A.** Schematic representation of the main lineage commitment steps in hematopoiesis, along with three immortalized cell lines (HPC7, G1E, G1E-ER4) and their approximate position relative to the primary cell populations shown. Abbreviations for cell populations are LSK = Lin-Sca1+Kit+ (which includes hematopoietic stem cells and multipotent progenitor cells), CMP = common myeloid progenitor cells, GMP = granulocyte monocyte progenitor cells, MEP = megakaryocyte erythrocyte progenitor cells, CLP = common lymphoid progenitor cells, CFUE = colony forming unit erythroid, ERY = erythroblasts, RBC = red blood cells, CFUMK = colony forming unit megakaryocyte, iMK = immature megakaryocytes, MK_fl = maturing megakaryocytes from fetal liver, PLTS = platelets, EOS = eosinophils, MAS = mast cells, NEU = neutrophils, MON = monocytes, T_CD8 = CD8+ T-cells, T_CD4 = CD4+ T-cells, B = B-cells, NK = natural killer cells. **B.** Available hematopoietic datasets. Shown in each row: Cell type along with its representative color, tissue stage (Ad = adult, ES diff = Embryonic stem cell derived, differentiated) and source (BM = bone marrow, sp = spleen, liver, blood). Shaded boxes indicate the presence of the dataset, and letters denote the source (V = VISION, L = Lara-Astiaso et. al 2014, O = other); see Supplemental Table S1 for more information. **C.** Correlations of nuclease accessible signals with all features (S3norm normalized) and across cell types. The genome-wide Pearson correlation coefficients *r* were computed for each cell type-feature pair and displayed as a heatmap after hierarchical clustering (using 1-*r* as the distance measure). The features are indicated by a characteristic color (first column on right), and the cell types are indicated in the second column to the right using the same colors as panel **B**. The full correlation matrix of all features across all cell types is in Supplemental Fig. S4.

The epigenomic data were gathered from many different sources, including individual laboratories and consortia (Fig. 1B and Supplemental Tables). These data had quality metrics within the ENCODE recommendations (see Supplemental Material section 2 and Supplemental Tables). However, this diversity of sources presented a challenge for data analysis, since each experiment differed widely in sequencing depth, fraction of reads on target, signal-to-noise ratio, presence of replicates, and other properties (Xiang et al. 2020), all of which can impact downstream analyses. We employed two strategies to improve the comparability of these heterogeneous datasets. First, the sequencing reads from each type of assay were uniformly processed, using pipelines similar to or adapted from current ENCODE pipelines (see Supplemental Material section 2). One notable difference is that our VISION pipelines allow reads to map to genes and genomic intervals that are present in multiple copies, thereby allowing interrogation of duplicated chromosomal segments, including multigene families and regions subject to deletions and amplifications. Second, for the ChIP-seq and nuclease accessibility data, we applied a new normalization method, S3norm, that simultaneously adjusts for differences in sequencing depths and signal-to-noise ratios in the collected data (Materials and Methods and Xiang et al. 2020). As with other normalization procedures, the S3norm method gives similar signals in common peaks for an epigenetic feature, but it does so without inflating the background signal. Preventing an increased background was necessary to avoid introducing spurious signals during the genome-wide modeling of the epigenetic landscape.

An overview of the similarities across all the datasets showed that most clustered by epigenetic features across cell types (Supplemental Fig. S4). For example, nuclease accessibility was highly correlated among the cell types examined, showing the global similarity in this primary feature of the regulatory landscape in blood cells (Fig. 1C). Other features such as CTCF and the signature marks for active promoters (H3K4me3) and enhancers (H3K27ac) showed notable but substantially lower correlations with the nuclease accessibility signal. In contrast, the H3K9me3 heterochromatin mark, the H3K27me3 Polycomb repressive mark, and the H3K36me3 had almost no correlation with nuclease sensitivity, and H3K4me1 showed modest correlation. The groupings within epigenetic features were more apparent after S3norm normalization (Supplemental Fig. S5), which supports the effectiveness of the normalization. The similarity of patterns for a particular feature across cell types suggested that combinations of features may be more effective than a single epigenetic mark to find patterns distinctive to a cell type.

In summary, our compilation of signal tracks, peak calls, estimates of transcript levels, and other material established a unified, consistently processed data resource for mouse hematopoiesis, which can be accessed at our VISION website (http://usevision.org).

### Simultaneous integration in two dimensions of non-binary epigenomic data

The frequent co-occurrence of some histone modifications has led to discrete models for epigenetic structures of candidate *cis*-regulatory elements, or cCREs (reviewed in Noonan and McCallion 2010; Hardison and Taylor 2012; Long et al. 2016). Moreover, the co-occurrences can be modeled formally using genome segmentation to learn the most frequently occurring, unique combinations of epigenetic features, called epigenetic states, and assigning each segment of DNA in each cell type to an epigenetic state. Computational tools such as chromHMM (Ernst and Kellis 2012), Segway (Hoffman et al. 2012), and Spectacle (Song and Chen 2015) provide informative segmentations primarily in one dimension, usually along chromosomes. The **I**ntegrative and **D**iscriminative **E**pigenome **A**nnotation **S**ystem (Zhang et al. 2016; Zhang and Hardison 2017), or IDEAS, expands the capability of segmentation tools in several ways. It integrates the data simultaneously in two dimensions, along chromosomes and across cell types, thus improving the precision of state assignments. It uses continuous (not binarized) data as the input, and the number of epigenetic states is determined automatically (Supplemental Fig. S6). Also, when confronted with missing data, it can make state assignments with good accuracy (Zhang and Mahony 2019).

When applied to the normalized epigenomic data from the 20 hematopoietic cell types, IDEAS learned 27 epigenetic states, including many expected ones as well as others that have been less frequently studied. The IDEAS model summary shows the prevalence of the eight epigenetic features in each state as a heatmap, organized by similarity among the states (Fig. 2A). The epigenetic state assignments were well supported by the underlying epigenomic data (Fig. 2B, Supplemental Fig. S3C). The epigenetic states described an informative landscape, distinguishing multiple state signatures representing distinct classes of regulatory elements (including enhancers, promoters and boundary elements). For example, six states showed a promoter-like signature, with high frequency of H3K4me3 (states 18, 21, 10, 15, 24, and 11); these are displayed in different shades of red, and P is the initial character in the explicit label. These six states distinguished promoter-like signatures by the presence or absence of other features with functional implications. For instance, the four promoter-like states that were also nuclease accessible (states 21, 10, 15, and 24) may encompass the nucleosome depleted region found adjacent to the transcriptional start site. Supporting this interpretation, three of these states (states 21, 10, and 24) also had the H3K27ac mark that frequently flanks the nucleosome-depleted region of active promoters. For all the major categories of chromatin associated with gene expression and regulation, including bivalent promoters, CTCF occupancy, enhancers, transcriptional elongation, repression, and heterochromatin, multiple states were discovered that differed in the combinations of associated features and their signal strengths. These are described in more detail in Supplemental Material section 7.

**Figure 2.**
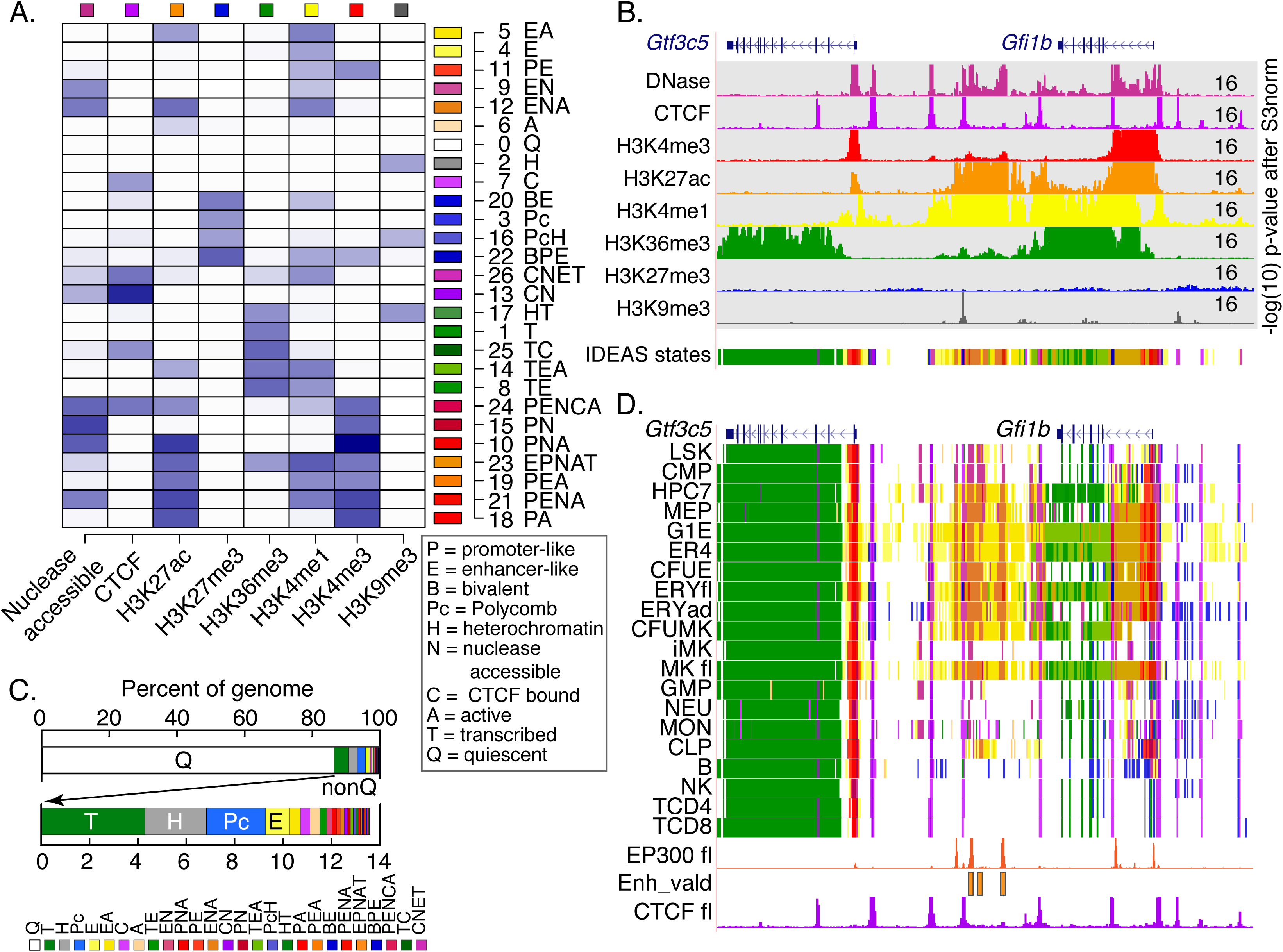
Segmentation of the epigenomes of hematopoietic cells after integrative modeling with IDEAS. **A.** Heatmap of the emission frequencies of each of the 27 states discovered by IDEAS, with state number and function-associated labels. Each letter in the label indicates a function associated with the combination of features in each state, defined in the *box*. The indicator for transcribed is H3K36me3, active is H3K27ac, enhancer-like is H3K4me1>H3K4me3, promoter-like is H3K4me3>H3K4me1, heterochromatin is H3K9me3, and polycomb is H3K27me3. **B.** Example of normalized epigenetic data from ERY in fetal liver around the *Gfi1b* locus, covering 70kb from chr2:28,565,001-28,635,000 in GRCm38/mm10, used as input to IDEAS for segmentation. The signal tracks are colored distinctively for each feature, with the inferred epigenetic states shown on the last track. The upper limit for signal in each normalized track is given at the right. **C.** Bar graphs of the average coverage of genomes by each state. The top graph emphasizes the high abundance of state Q, and the second graph shows the abundances of the 26 non-quiescent states. The key for annotated colors is the same order as the states in the bar graph. **D.** Segmentation pattern across cell types around the *Gfi1b* exemplar locus. Signal tracks for EP300 (ENCSR982LJQ, ENCODE consortium) and CTCF from mouse fetal liver were included for validation and confirmation, along with the locations of enhancers shown to be active (Enh_vald; Moignard et al. 2013).

The fraction of the genome in each state reveals the proportion of a genome associated with a particular activity. The most common state in all the epigenomes is quiescence, i.e. state 0 with low signals for all the features (Fig. 2C). The mean percentage of the genome in this state was 86%, with values ranging from 85% to 92% in individual cell types. About 60% of the genome was in this state in all cell types examined, indicating that in hematopoietic cells, about 40% of the mouse genome is incorporated within chromatin with the dynamic histone modifications identified in this study. The most common non-quiescent states were transcribed, heterochromatic, and Polycomb repressed (Fig. 2C). The remaining portion of the genome was populated with a large number of active states, comprising ~4% of the genome. Thus, only a small proportion of the genome in each cell type was found in chromatin associated with the dynamic histone modifications assayed here. This small fraction of the genome is probably responsible for much of the regulated gene expression characteristic of each cell type.

### Visualizing the regulatory landscape across hematopoietic cell types as defined by the IDEAS segmentation

The chromatin activity landscape inferred by IDEAS can be displayed by assigning the distinctive color for each state to DNA segments along chromosomes and across cell types. (Fig. 2D). For example, genes transcribed in all cell types, such as *Gtf3c5*, were painted red at the active promoter and green for regions of transcriptional elongation. Within and between the transcription units were short purple segments indicating CTCF binding, aligning with the CTCF occupancy data available for tissues like fetal liver and providing a prediction for CTCF binding in other cell types. The gene *Gfi1b*, encoding a transcription factor required in specific hematopoietic lineages, showed different state assignments across the cell types, with active promoters (red), intronic enhancers (orange), and transcribed regions (green) in CMP, erythroid, and megakaryocytic cells but fewer active states in other cell types. Downstream (left) of *Gfi1b* was a large region with many DNA segments assigned to enhancer-associated states; these were model-generated candidates for regulating expression of *Gfi1b*. The potential roles of the intronic and downstream candidate enhancers were supported by binding of the coactivator EP300 observed both in mouse fetal liver and MEL cells (Yue et al. 2014; The ENCODE Project Consortium et al. 2020), information that was not included in training the model. Furthermore, previous studies of cross-regulation between GATA2 and GFI1B revealed three enhancers downstream of the *Gfi1b* gene by reporter gene assays in transgenic mouse and transfected cells (Moignard et al. 2013). These enhancers overlapped with the model-predicted enhancers and provided strong experimental validation of the predictions from the IDEAS segmentation.

### cCREs across mouse hematopoiesis

While genomic regions potentially involved in gene regulation can be discerned from the segmentation views of regulatory landscapes, it is important to assign discrete genomic intervals as candidate *cis*-regulatory elements (cCREs) to clarify assessments and validations of regulatory elements and to empower systematic modeling of regulatory systems. Therefore, we combined our nuclease sensitivity data with IDEAS segmentation to infer a set of 205,019 cCREs in the 20 cell types.

A cCRE was defined as a DNA segment assigned as a reproducible peak by ATAC-seq or DNase-seq that was not in a quiescent epigenetic state in all cell types (Supplemental Fig. S8). We considered ATAC-seq or DNase-seq data to be reproducible when peaks were called in each replicate (when replicates were available). Some peaks were assigned to the quiescent state in all cell types, and these were removed from the set of cCREs. No cell type-specific cCREs could be called in mature MK or CLP cells because no ATAC-seq or DNase-seq data were available for these cell types; however, we inferred the epigenetic states in these two cell types for the DNA segments predicted to be cCREs in other cell types. This information about the locations and epigenetic states of cCREs in hematopoietic cell types provides a valuable resource for detailed studies of regulation both at individual loci and globally across the genome.

Because a wide range of hematopoietic cells was interrogated for epigenetic features, we expected that the set of cCREs from the VISION project would expand and enhance other collections of cCREs. Thus, we compared the VISION cCRE set with the Blood Cell Enhancer Catalog, which contains 48,396 candidate enhancers based on iChIP data in sixteen mouse hematopoietic cell types (Lara-Astiaso et al. 2014), and a set of 56,467 cCREs from mouse fetal liver released by the ENCODE project (The ENCODE Project Consortium et al. 2020). Furthermore, we examined the set of 431,202 cCREs across all assayed mouse tissues and cell types in the SCREEN cCRE catalog from ENCODE (The ENCODE Project Consortium et al. 2020). The overlapping DNA intervals among combinations of datasets revealed substantial consistency in the inferred cCREs (Fig. 3A). A large proportion of the VISION cCREs (70,445 or 41.5%) were in the iChIP Blood Enhancer Catalog and/or the SCREEN fetal liver cCREs. Conversely, a majority of the cCREs in the iChIP catalog (78.7%) were also in VISION cCREs, as expected given the large contribution of iChIP data to the VISION compilation. An even larger proportion (84%) of the SCREEN fetal liver catalog was in VISION cCREs. The cCREs that are common among these collections, despite differences in data input and analysis, are strongly supported as candidate regulatory elements.

**Figure 3.**
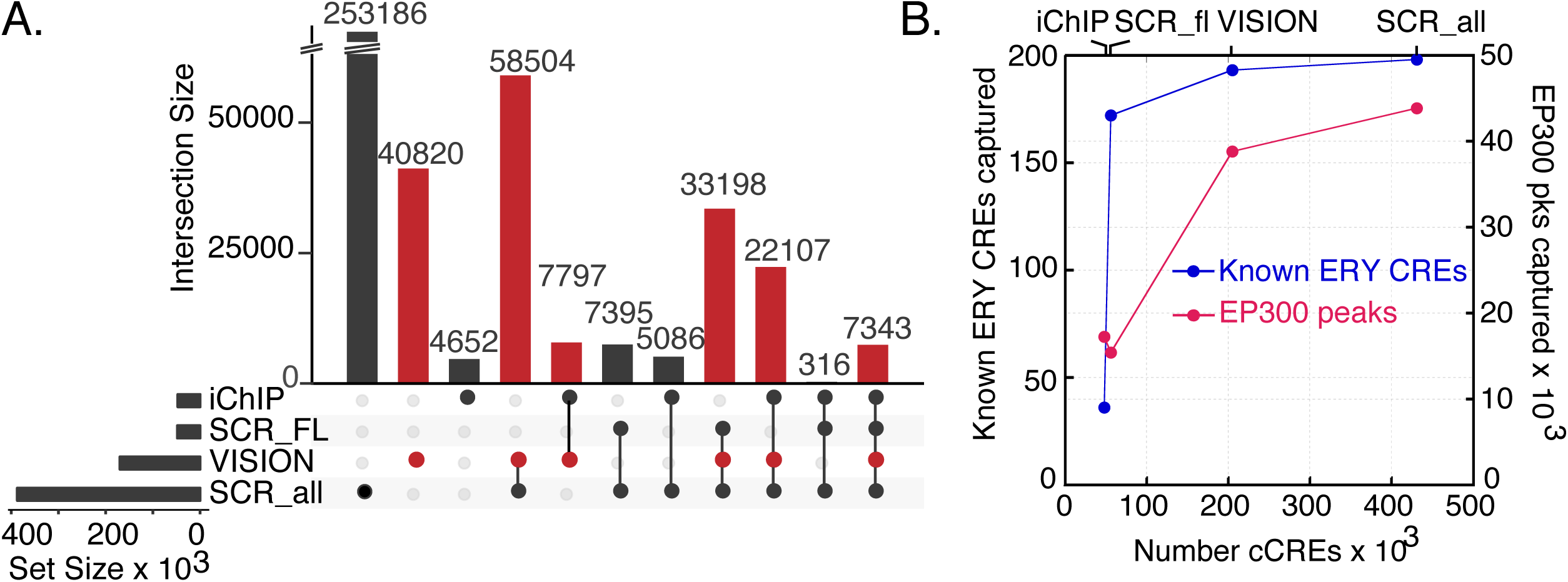
Comparative analysis of VISION cCREs. **A.** Overlaps of the VISION cCREs with three other cCRE catalogs. The overlapping cCREs in all four datasets were merged. The numbers of merged cCREs in each set were labeled on each row, and the numbers in each level of overlap were shown in columns, visualized using an UpSet plot (Lex et al. 2014). The sets compared with the VISION cCREs were the Blood Enhancer Catalog derived from iChIP data (iChIP; Lara-Astiaso et al. 2014), the SCREEN cCREs specific to mouse fetal liver at E14.5 (SCR_FL), and those for all tissues and cell types in mouse (SCR_all). **B.** The VISION cCREs capture known regulatory elements and orthogonal predicted cCREs. The number of known CREs that are also present in each cCRE collection was plotted against the number of regulatory elements (known or inferred) in each dataset. The EP300 peaks were deduced from EP300 ChIP-seq data from ENCODE, reprocessed by VISION pipelines, from FL E14.5, MEL, and CH12 cells. Replicated peaks were combined into one dataset and merged, to get over 60,000 peaks. The number of known EP300 peaks that were also present in each cCRE collection was plotted against the number of cCREs in each dataset.

The VISION cCRE set is substantially larger than either the iChIP Blood Enhancer Catalog or the SCREEN fetal liver cCREs, and we hypothesized that the larger size reflected the inclusion of greater numbers of cell types and features in the VISION catalog. This hypothesis predicts that VISION cCREs that were not in the other blood cell cCRE sets may be found in larger collections of cCREs, and we tested this prediction by comparing VISION cCREs to the entire set of ENCODE SCREEN cCREs. Indeed, we found another 58,504 (34.5%) VISION cCREs matching this catalog across mouse tissues, supporting the interpretation that the VISION cCRE set is more comprehensive than other current blood cell cCRE collections. Overall, the comparisons with other collections supported the specificity and accuracy of the VISION cCRE set.

To further assess the quality of the VISION cCRE set, we evaluated its ability to capture known *cis*-regulatory elements (CREs) and independently determined DNA elements associated with gene regulation. Using a collection of 212 experimentally determined, erythroid CREs curated from the literature (Dogan et al. 2015) as known erythroid CREs, we found that while the iChIP Blood Enhancer catalog captured only a small portion, the VISION and SCREEN fetal liver cCREs overlapped with almost all the erythroid CREs (Fig. 3B). The latter two collections were built from datasets that included highly erythroid tissues, such as fetal liver, which may explain their more complete coverage than the Blood Enhancer Catalog, which was built from datasets from fewer erythroid cell types. Increasing the number of cCREs to over 400,000 in the SCREEN mouse cCREs did not substantially increase the number of known CREs that overlap. Thus, the VISION cCREs efficiently captured known erythroid CREs.

The co-activator EP300 catalyzes the acetylation of histone H3K27, and it is associated with many active enhancers. We used ChIP-seq data on EP300 as a comparison set of blood cell candidate enhancers that were determined independently of the data analyzed in VISION. The ENCODE consortium has released replicated datasets of EP300 ChIP-seq data determined in three blood-related cell types from mouse, MEL cells representing maturing proerythroblasts, CH12 cells representing B cells, and mouse fetal liver from day E14.5 (Yue et al. 2014; The ENCODE Project Consortium et al. 2020). After re-processing the ChIP-seq data using the VISION project pipelines, replicated peaks were merged across the cell types to generate a set of over 60,000 EP300 peaks in blood related cells. The VISION cCRE set efficiently captured the EP300 peaks, hitting almost two-thirds of these proxies for regulatory elements, a much larger fraction than captured by the Blood Enhancer catalog or ENCODE fetal liver cCREs (Fig. 3B). Expanding the number of SCREEN cCREs to over 400,000 gave only a small increase in the number of EP300 peaks captured. The EP300 peaks not captured by the VISION cCREs tended to have lower signal strength and were less associated with ontology terms such as those for mouse phenotype (Supplemental Fig. S9), suggesting that VISION cCREs captured the more likely functional EP300 peaks.

These analyses show that the VISION cCREs included almost all known erythroid CREs and they captured a large fraction of potential enhancers identified in relevant cell types by a different feature (EP300).

### Global comparisons of regulatory landscapes and transcriptomes

The collection of cCREs and transcriptomes in VISION provided an opportunity to examine the relationships between cell types, including both purified populations of primary cells and cell lines. In conducting this analysis, we distinguished a cCRE from an active cCRE. A cCRE, which is a DNA interval predicted to be a regulatory element in any cell type, is present in all cell types, just as a gene is present in all cell types. However, a cCRE can show evidence of activity (either positive or negative) differentially across cell types, just as genes may be active in only some cell types. Thus, we refer to cCREs in epigenetic states indicative of regulatory activity as active cCREs, including states with either positive or negative associations with gene expression.

The epigenetic modifications at cCREs are a prominent feature of the regulatory landscape. Thus, to compare the regulatory landscape across cell types, we used the correlations between the nuclease accessibility signals in cCREs across cell types to group the cell types by hierarchical clustering (Fig. 4A). All erythroid cell types, including the G1E and G1E-ER4 cell lines, clustered with MEP to the exclusion of other cell types. The remaining cell types formed two groups. One consisted of hematopoietic stem and multilineage progenitor cells (LSK, CMP and GMP) along with early progenitor (CFUMK) and immature (iMK) megakaryocytic cells. The other contained both innate (NEU, MON) and acquired (B, NK, T-CD4, T-CD8) immunity cells. Comparisons using a dimensional reduction approach (principal component analysis or PCA) also supported these groupings (Supplemental Fig. S10A).

**Figure 4.**
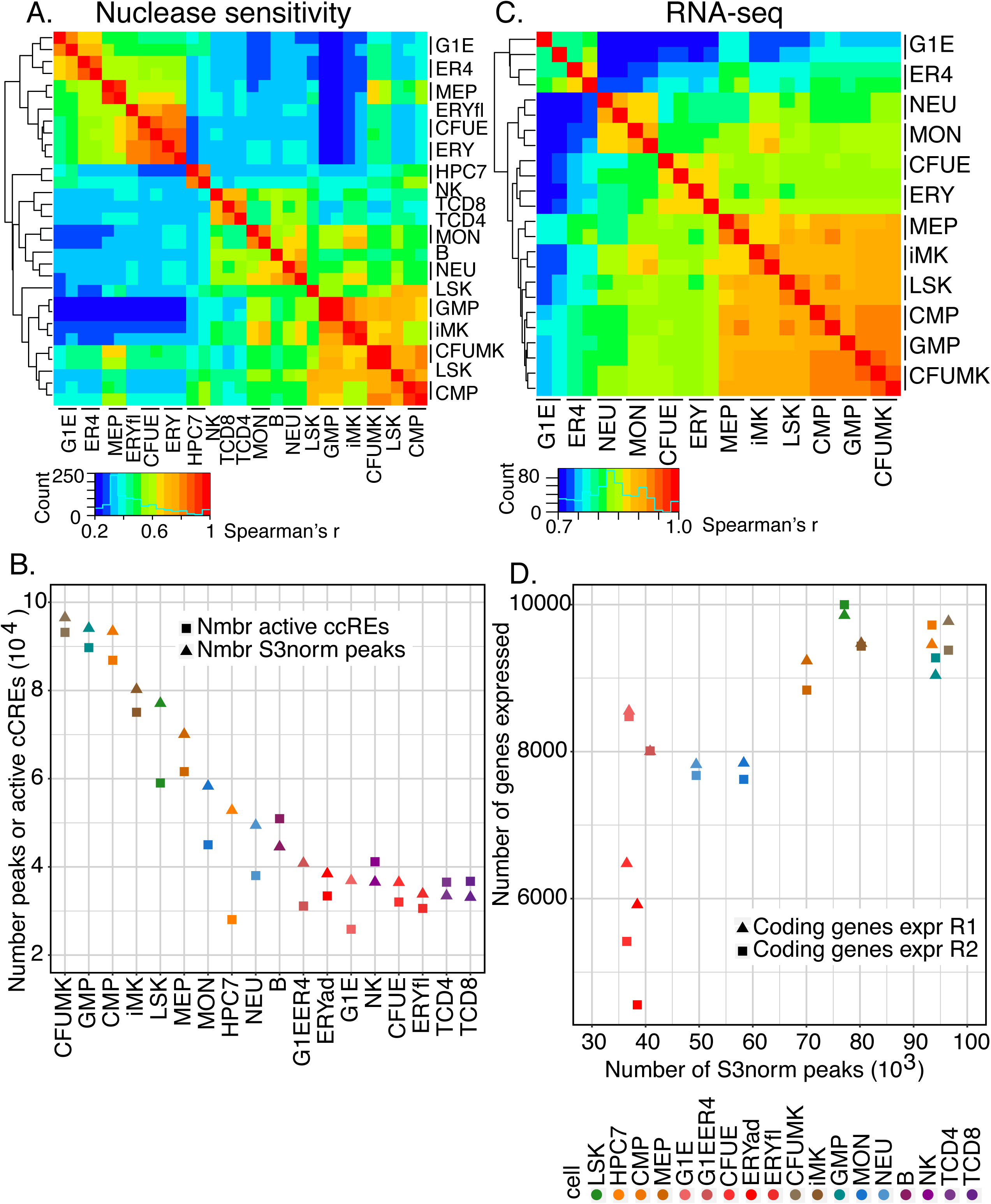
Global comparisons of nuclease accessibility profiles and transcriptomes across mouse hematopoietic cell types. **A.** Heatmap of the hierarchical clustering of nuclease-sensitive elements (ATAC-seq and DNase-seq, using S3norm for normalization), with Spearman’s rank correlation *r* as the similarity measure, and 1-*r* as the distance measure for hierarchical clustering across 18 cell types. Results include replicates for cell types with replicated data (indicated by bars next to the cell type name). **B.** Numbers of dynamic cCREs in each cell type, determined from ATAC-seq and DNase-seq profiles, and analyzed as both peak calls from Homer or from peaks after S3 normalization. **C.** Heatmap of the hierarchical clustering of RNA-seq (TPM values for all genes, quantile normalized, showing replicates), with Spearman’s *r* as the similarity measure. **D.** Concordant decreases during hematopoietic differentiation in nuclease accessibility and expressed genes, shown as the association between numbers of genes expressed and numbers of dynamic cCREs across cell populations and types.

Furthermore, the PCA and subsequent analyses showed that a substantial reduction in the number of active cCREs was a major contributor to the differences in the landscape of nuclease accessibility during hematopoietic differentiation. The first principal component (PC1) captured a large fraction (82%) of the variation, placing the cell types along an axis with many multilineage progenitor cells on one end and many mature cells on the other (Supplemental Fig. S10A). That PC1 axis was highly correlated with the numbers of active cCREs (Supplemental Fig. S10B, and a direct comparison showed a progressive decline in numbers of cCREs active in most maturing blood cells (Fig. 4B). We conclude that a reduction in numbers of active cCREs is a major trend during mouse hematopoietic differentiation.

The gene expression landscape was also compared across cell types, using estimates of gene transcript levels from RNA-seq data in a subset of 12 cell types interrogated by the same method within our VISION laboratories. RNA-seq data on acquired immunity cells were not included because the assay was done by a substantially different procedure (Lara-Astiaso et al. 2014), and this difference in RNA-seq methodology dominated the combined comparison. The hierarchical clustering results (Fig. 4C) and PCA (Supplemental Fig. S10C) revealed three clusters that were largely consistent with the analysis of the regulatory landscape, grouping megakaryocytic cells with multilineage progenitors while keeping primary erythroid cells (CFUE and ERY) and innate immune cells (NEU and MON) in distinct groups. In contrast, MEP cells grouped with progenitor cells in the transcriptome profiles whereas they grouped with erythroid cells by nuclease sensitivity data. MEP cells have a pronounced erythroid bias in differentiation (Psaila et al. 2016), and this difference in the grouping of MEPs suggests that the regulatory landscape of MEP has shifted toward the erythroid lineage prior to reflecting that bias in the transcriptome data. G1E and G1E-ER4 cell lines, which are models for GATA1-dependent erythroid differentiation, also were placed differently based on cCRE and transcriptome data, forming a separate cluster in the transcriptome data. While that result reveals a difference in the overall RNA profiles between G1E and G1E-ER4 cells versus primary cells, their grouping with primary erythroid cells by cCRE landscape supports the use of these cell lines in specific studies of erythroid differentiation.

The decrease in numbers of cCREs during differentiation and maturation was associated with a decrease in numbers of genes expressed. The highest numbers of protein-coding genes were expressed in the progenitor (LSK, CMP, GMP, MEP) and megakaryocytic (CFUMK and iMK) cells, with fewer in MON and NEU, and the lowest number in erythroid cells (CFUE and ERY) (Fig. 4D). A larger number of genes were expressed in the ES-derived cell lines, G1E and G1E-ER4, than in the primary erythroid cells. A similar decline was observed over a ten-fold range of thresholds for declaring a gene as expressed (TPM exceeding 1, 5 or 10). The parallel decreases in numbers of active cCREs and expressed genes led to a strong positive association between these two features (coding genes: Fig. 4D; Pearson correlation *r*= 0.90 or 0.78 when values for G1E and G1E-ER4 cells were excluded and included, respectively, in a linear fit; noncoding genes: Supplemental Fig. S10D). Similar results were reported for transitions during megakaryopoiesis and erythropoiesis in Heuston et al (2018) based on peak calls for histone modification and nuclease accessibility. Our results based on integrative modeling confirm these conclusions and show that the reduction in numbers of expressed genes and active cCREs was observed broadly across hematopoiesis. Considering specifically genes encoding hematopoietic regulators, we found that this general decline in transcription led to a reduction in the number of hematopoietic regulators produced in differentiated, maturing erythroid cells but not in other hematopoietic cell types (Supplemental Fig. S11). We conclude that the breadth of transcription declines during differentiation, and furthermore the loss of activity of cCREs may contribute to the decrease in numbers of genes expressed.

### Epigenetic states of cCREs vary across cell types in an informative manner

The VISION catalog of cCREs, annotated by their epigenetic state in each cell type, can be used to track both the timing and types of transitions in epigenetic states during differentiation, which provide insights into regulatory mechanisms, e.g. which CREs are likely to be inducing or repressing a target gene. The full scope of state transitions in cCREs across cell types is complex, and in this section, we focus on major transitions contributing to changes in the numbers and state of active cCREs.

Within the dominant pattern of decreasing numbers of active cCREs during commitment and maturation of lineages (except MK), the reduction was particularly pronounced for cCREs in state 9 (EN) and state 13 (CN) (Fig. 5A), while changes in the numbers of cCREs in other states were more modest (Fig. 5B, Supplemental Fig. S12 A, B). These state-specific reductions suggested that many active cCREs in progenitor and MK cells were in a poised enhancer mode (state 9 EN) or in a CTCF-bound, nuclease accessible state (state 13 CN). We then determined the states into which these cCREs tended to transition by examining all state transitions in cCREs between all pairs of cells. In the case of CMP cells differentiating to ERY, we found that cCREs in the poised enhancer state 9 in CMP did not tend to stay in state 9, but rather they more frequently transitioned to states 12 (active enhancer), 3 (polycomb), and 0 (quiescent) in ERY (Supplemental Fig. S12C). These classes of state transitions were strongly supported by examination of the underlying signals for the nuclease sensitivity and histone modifications (Fig. 5C). Discrete classification of cCREs by their state assignments across cell types also reveal these major transitions (Supplemental Fig. S13). This systematic analysis of transitions in epigenetic states across cell types helps uncover the differentiation history of cCREs and provides mechanistic insights into regulation. For example, using SeqUnwinder (Kakumanu et al. 2017) to discover discriminative motifs, we found that the CMP cCREs that transition from poised to active enhancer in the erythroid lineage were enriched for the GATA transcription factor binding site motif, whereas those that transition to a polycomb state were enriched in motifs for binding ETS transcription factors such as PU.1 (Supplemental Fig. S14). These results are consistent with the known antagonism between GATA1 and PU.1 in erythroid versus myeloid differentiation (Rekhtman et al. 1999; Zhang et al. 1999). Thus, they illustrate the value of machine-learning approaches, such as assigning epigenetic states systematically and finding discriminative motifs, to uncover relationships from genome-wide data that fit with models derived from decades of experimentation.

**Figure 5.**
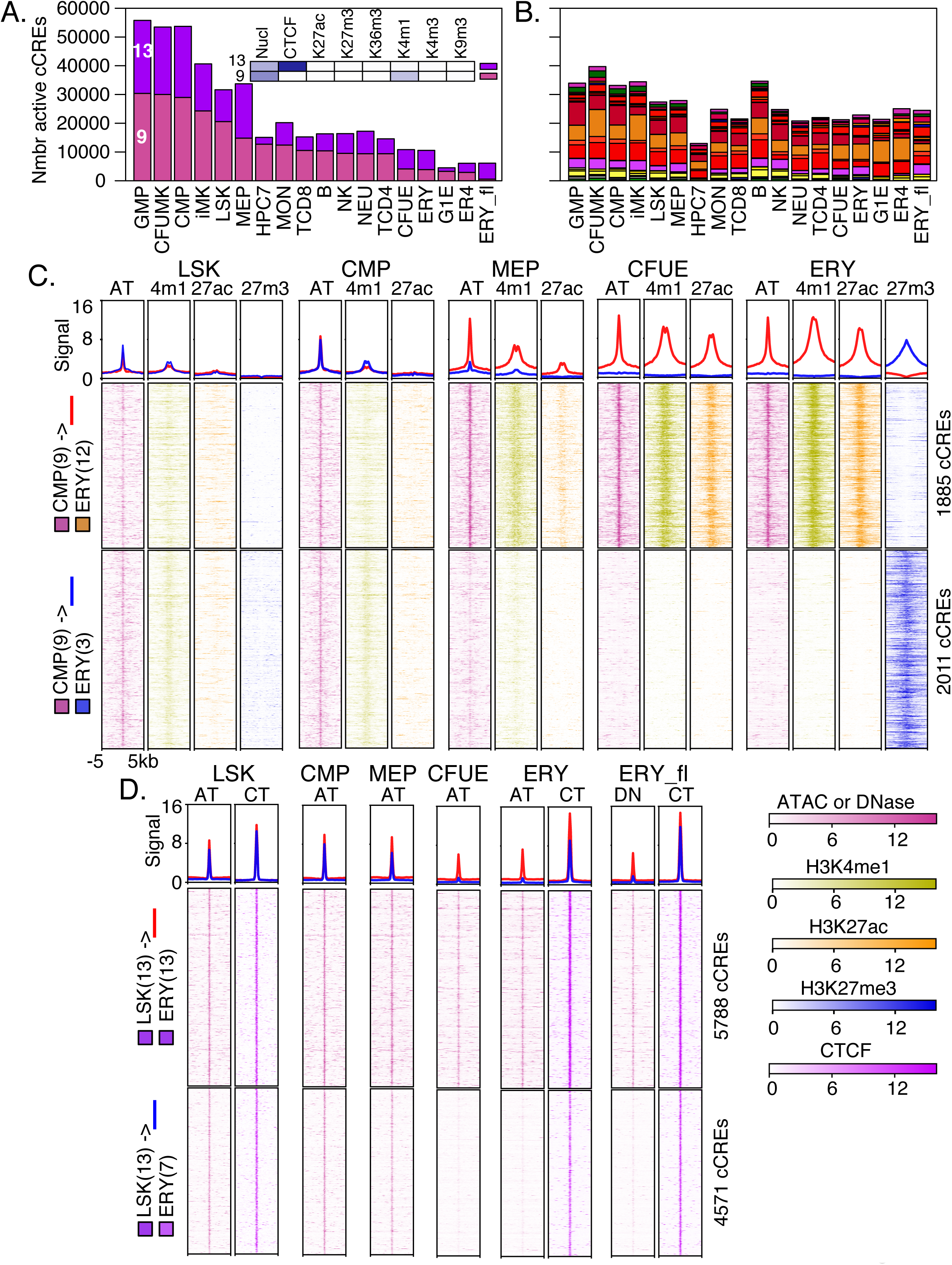
Transitions in epigenetic states at cCREs across hematopoietic differentiation. **A and B.** The numbers of cCREs in each cell type are colored by their IDEAS epigenetic state, emphasizing decreases in numbers of cCREs in states 9 and 13 (**A)**, while numbers in other states less variable (**B**). **C**. Aggregated and individual signal profiles for cCREs in the poised enhancer state 9 in CMPs as they transition from LSK through CMP and MEP to CFUE and ERY. Profiles for up to four relevant epigenetic features are presented. Data for H3K27me3 are not available for CMP, MEP, or CFUE. The first graph in each panel shows the aggregated signal for all cCREs in a class, and graphs beneath it are heatmaps representing signal intensity in individual cCREs. In the aggregated signal, red lines show signals for cCREs that transition from poised state 9 to active enhancer-like state 12, and blue lines show signals for cCREs that transition from poised state 9 to polycomb repressed state 3. **D**. Aggregated and individual signal profiles for CTCF-bound cCREs that either retain or lose nuclease accessibility during differentiation from LSK to ERY. In the aggregated signal, red lines show signals for cCREs that stay in the CTCF-bound, nuclease sensitive state 13, and blue lines show signals for CTCF-bound cCREs that lose nuclease sensitivity as they transition from state 13 to state 7. Signals were normalized by S3norm. Abbreviations are AT=ATAC, 4m1=H3K4me1, 27ac=H3K27ac, 27m3=H3K27me3, CT=CTCF.

Another major state of cCREs in progenitor and megakaryocytic cells was CTCF-bound and nuclease accessible (state 13). Much of the decrease in numbers of cCREs in this state occurred through a loss of accessibility while retaining occupancy by CTCF (state 7, Supplemental Fig. S12C and D). To eliminate the possibility that the inferred loss of nuclease sensitivity was an artifact of low sensitivity in the ATAC-seq data, we examined these cCREs for DNase sensitivity in an independent experiment conducted on ERY from fetal liver (ERY_fl). We found that the cCREs undergoing the transition from state 13 to state 7 had low nuclease sensitivity in ERY by both assays, as well as in CFUE, while retaining a strong CTCF signal (Fig. 5D). Thus, we concluded that the state 13 to state 7 transition was not an artifact of poor sensitivity of the accessibility assays. The loss of nuclease accessibility at this subset of CTCF-bound sites occurred between MEP and CFUE stages, suggesting that it could be connected to the process of erythroid commitment. By examining genes in the vicinity of the CTCF-bound cCREs, we found that this loss of nuclease sensitivity at CTCF-bound sites occurred in more gene-poor regions, and it was associated to some extent with gene repression (Supplemental Figure S15). The CTCF-bound cCREs that retained nuclease accessibility during differentiation were enriched at TAD boundaries that were common across myelo-erythroid differentiation (Supplemental Fig. S16).

In summary, the number of active cCREs declined dramatically as cells differentiated from stem and progenitor cells to committed, maturing blood cells. This decrease in cCREs was strongly associated with a reduction in the numbers of expressed genes in committed cells. Our analysis of epigenetic states in cCREs across this process revealed major declines in two states. First, the poised enhancer state was prevalent in cCREs in stem and progenitor cells, and it had two major fates. One was a transition to an active enhancer state, and in the erythroid lineage this transition was associated with GATA transcription factor binding site motifs, as expected for activation of erythroid genes. The other fate was to lose nuclease sensitivity and switch to a repressed state. Those state transitions were not novel observations, but our extensive annotation of the cCREs allows investigators to identify which cCREs around genes of interest make those transitions. Second, another state prevalent in stem and progenitor cells was a CTCF-bound and nuclease accessible state. The number of cCREs in that state declined during differentiation, with many cCREs transitioning to a state with CTCF still bound but no longer nuclease accessible. Further studies are needed to better understand the roles of these different classes of CTCF-bound sites.

### Estimating regulatory output and assigning target genes to cCREs

We investigated the effectiveness of the collection of mouse hematopoietic cCREs from VISION in explaining levels of gene expression. We developed a modeling approach to evaluate how well the cCREs, in conjunction with promoters, could account for levels of expression in the twelve cell types for which the RNA-seq measurements were determined in the same manner. This modeling approach had the additional benefit of making predictions of target genes for each cCRE.

We reasoned that the epigenetic state assignments for each cCRE DNA interval in each cell type could serve as a versatile proxy for cCRE regulatory activity, since the states were based on a systematic integration of multiple epigenetic signals. As explained in detail in the Supplemental Material, section 17, we estimated promoter and cCRE effects on expression by treating the states as categorical variables and training a multivariate linear model of gene expression on the states. Each cCRE and promoter could be composed of multiple epigenetic states (Fig. 6A), and we used the proportion of promoters and the proportion of pooled cCREs covered by a state as the predictor variable for that state (Fig. 6B). However, in our sub-selection training, a given cCRE is represented by a single state rather than a weighted sum of states (Supplemental Material). All cCREs within 1Mb of the TSS of a gene were initially considered and then filtered by a minimum correlation to that gene’s expression. Not all cCREs within the 2 Mb region surrounding a gene’s TSS were expected to influence expression. Thus, CREs predicted to have limited contribution to explaining expression were removed via a sub-selection strategy during iterations of model fitting (Fig. 6B, Supplemental Fig. 17B).

**Figure 6.**
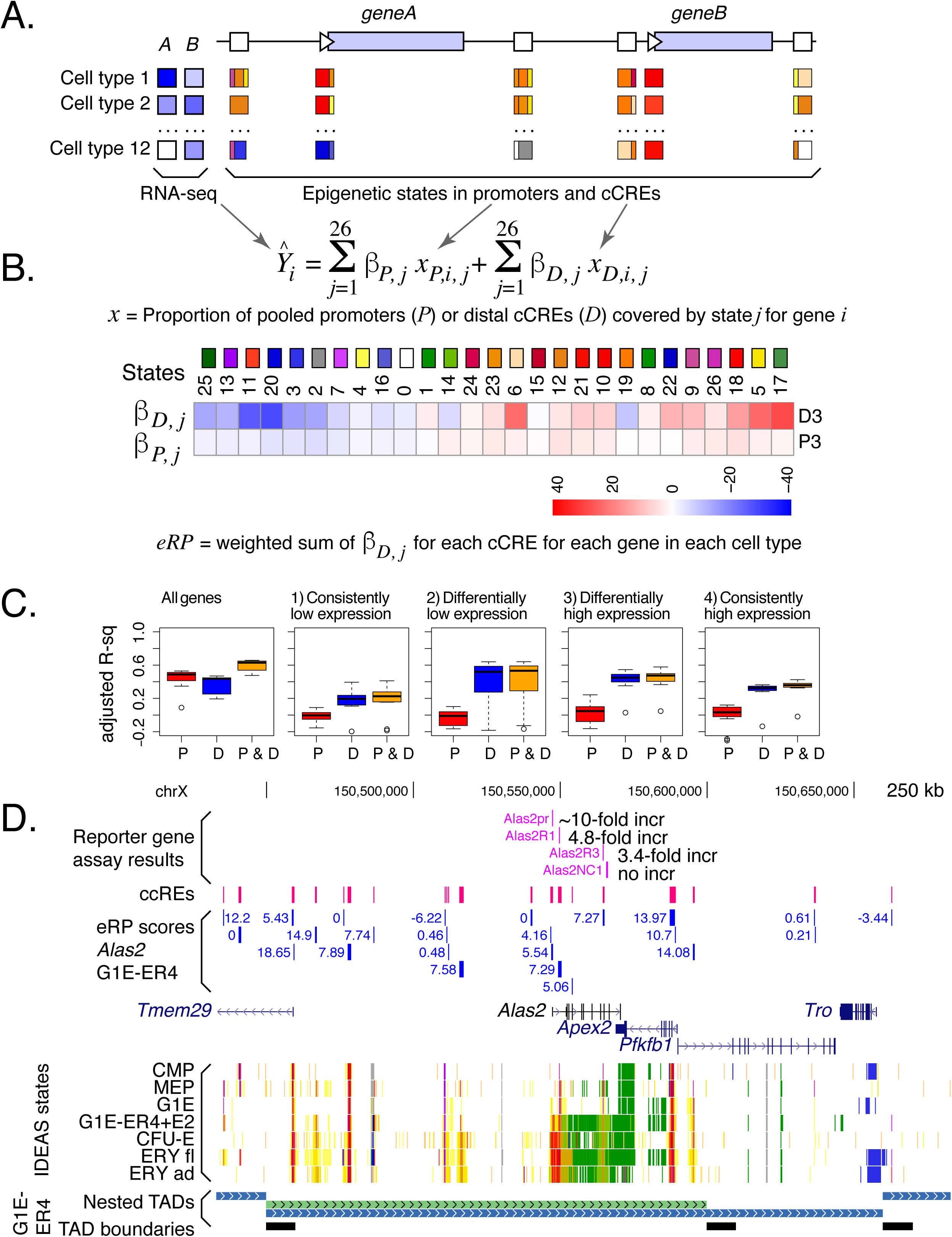
Initial estimates of regulatory output and target gene prediction using regression models of IDEAS states in promoters and cCREs versus gene expression. **A.** Illustration of promoters and cCREs around two potential target genes, showing expression profiles of the genes across cell types (shades of blue, *left*) and promoters/cCREs with one or more epigenetic states assigned in each cell type. **B.** Multivariate linear regression of proportion of promoters and pooled cCREs in each state against expression levels of potential target genes, keeping promoters and cCREs separate and learning the regression coefficients iteratively in a sub-selection strategy. Values of the regression coefficients beta for each epigenetic state for promoters and cCREs for differentially expressed genes. The values of the regression coefficients for each epigenetic state are presented as a blue to red heatmap. **C.** Ability of eRP scores of cCREs to explain levels of expression on chr1-chr19 and chrX in the twelve cell types for all genes and (1-4) in the four categories of genes. A leave-one-out strategy was employed to calculate the accuracy predicting expression. The distribution of adjusted *r*^2^ values are shown as box-plots for promoters, distal cCREs, and combined. **D.** Illustration of eRP scores for cCREs in and around the *Alas2* gene, including a comparison with previously measured enhancer and promoter activities. Nested TADs called by OnTAD (An et al. 2019) are shown in the bottom tracks.

The regression coefficients, *β*, determined for the epigenetic states showed some expected trends. For example, the coefficients for the set of differentially expressed genes were high for most promoter-like and enhancer-like states and low for most polycomb-repressed and heterochromatin states (Fig. 6B, and a full set of values is presented in Supplemental Fig. 17D).

We evaluated the accuracy of predicting gene expression from the weighted sum of the state-specific regression coefficients using a leave-one-out strategy. Specifically, we trained a linear model on data from eleven of the twelve cell types, minimizing mean squared error (MSE), and then computed the adjusted *r*^2^ for the accuracy of the predicted expression levels compared to the actual expression levels in the left-out cell type. This procedure was repeated leaving out each of the cell types in turn. Coefficients were calculated using only promoters, only cCREs, or a combination of both. In the case of the cCRE trained model, we defined the sum of coefficients weighted by each cCRE state proportion as the epigenetic regulatory potential (eRP) score. The predicted expression for each gene was the mean of the eRP scores for all paired cCREs. For expression of all genes, the prediction accuracy was around 50% for promoters only or eRPs only, and it improved to about 60% when both were combined (Fig. 6C, graph All genes).

Some portion of the explanatory power was expected to derive from the strong differences in epigenetic signals for expressed *versus* silent genes. In an effort to remove this effect from the predictions of accuracy, we repeated the linear regression modeling and evaluations on four categories of genes separately, specifically those with (1) consistently low, (2) differentially low, (3) differentially high, and (4) consistently high expression across cell types. The values of *β* varied across the four categories (Supplemental Fig. 17D). Using gene category partitioning, the accuracy of predicting expression levels in the leave-one-out strategy showed a much smaller impact of the promoters (Fig. 6C, graphs 1-4), suggesting that a major effect of the epigenetic states around the TSSs was to establish expression or silencing. In contrast, the distal cCREs did contribute to expression variation within the gene categories, especially for differentially expressed genes (Fig. 6C, graphs 2 and 3). Overall, these evaluations indicate that promoters contributed strongly to the broad expression category (expressed or not, differential or constitutive), and distal cCREs contributed to the expression level of each gene within a category.

By considering these linear regression coefficients as proxies for the regulatory output of cCREs in a particular epigenetic state, we used them to estimate the impact of histone modifications around cCREs close to differentially expressed genes. Many expected associations were found, but in addition, this analysis revealed that H3K27ac was the histone modification at cCREs most distinctly associated with gene activation, CTCF at a cCRE was associated with repression, and H3K4me1 and nuclease accessibility were about equally frequent in states with positive or negative impacts on expression (Supplemental Fig. S18).

The positive predictive power of these initial estimates of eRP scores supported their utility in assigning candidates for target genes for cCREs. The estimated eRP scores can serve as one indicator of the potential contribution of each cCRE to the regulation of a gene in its broad vicinity. Thus, a set of likely cCRE-target gene pairs can be obtained at any desired eRP threshold. We provide a large table of potential cCRE-target gene pairs at the VISION project website, along with a versatile filtering tool for finding cCREs potentially regulating a specified gene in a particular cell type. The filtering tool also allows further restriction of cCREs to those within the same topological associated domain or compartment as the candidate target gene. The example from the *Alas2* locus (Fig. 6D) illustrates how these eRP scores were consistent with results from previous experimental assays for CREs within the gene (Wang et al. 2006), and they raise the possibility of additional, distal cCREs regulating the gene. These data-driven, integrated resources should allow users to make informed decisions about important but challenging issues such as finding the set of cCREs likely to regulate a particular gene.

## Discussion

One goal of the VISION project is to gather information from our laboratories, other laboratories, and consortia to conduct systematic integrative analysis and produce resources of high utility to investigators of genome biology, blood cell differentiation, and other processes. In this study, we compiled and generated epigenomic and transcriptomic data on cell types across hematopoietic differentiation in mouse. The data were systematically analyzed by the IDEAS method to assign genomic intervals to epigenetic states in twenty cell types, with each state defined by a quantitative spectrum of nuclease sensitivity, histone modifications, and CTCF occupancy. Most of these combinations of epigenetic features are associated with specific regulatory elements or events, such as active promoters, poised enhancers, transcribed regions, or quiescent zones, and thus, the epigenetic state assignments provide a guide to potential functions of each genomic interval in each cell type. In effect, the IDEAS segmentation pipeline reduced 150 dimensions (or tracks) of epigenomic data to twenty dimensions, i.e. the number of cell types examined. While the cell populations studied can be conceptualized as cell “types”, it is important to keep in mind that these populations, especially of stem and progenitor cells, are heterogeneous, and thus our integrative analyses do not delve into all the stages of hematopoietic differentiation and maturation. We further focused the epigenomic data by constructing an initial registry of 205,019 cCREs, which are discrete genomic intervals with features predictive of a potential regulatory role in one or more hematopoietic cell types, along with state assignments and initial estimates of regulatory output for candidate target genes in each cell type. Investigators now have simplified ways to view the large amount of data, e.g. in a genome browser, and to operate computationally on the state assignments and cCREs.

We provide multiple ways for investigators to access and interact with the data via our VISION website (usevision.org). The raw and normalized data tracks can be downloaded for further analysis. The regulatory and transcriptomic landscapes around individual genes can be viewed in our custom genome browser, which is built on the familiar framework of the UCSC Genome Browser (Haeussler et al. 2019). Tables of annotated cCREs and their associations with specific genes by regression can be downloaded, and cCREs for specific genes and genomic intervals can be obtained by queries at the website. Links are provided to additional resources such as CODEX for more extensive transcription factor occupancy and histone modification data (Sanchez-Castillo et al. 2015), the 3D Genome Browser for visualizing matrices of chromatin interaction frequencies (Wang et al. 2018), and the ENCODE registry of cCREs (The ENCODE Project Consortium et al. 2020).

We chose IDEAS as the systematic integration method because its joint segmentation along chromosomes and across cell types retains position-specific information, thereby providing more precision to the state assignments (Zhang et al. 2016; Zhang and Hardison 2017). Furthermore, the IDEAS method does not require determination of all features in all cell types, and thus cell types with missing data were included (Zhang and Mahony 2019). Even an extreme case of the cell type CFUMK, for which the only epigenomic dataset was ATAC-seq, was assigned a meaningful segmentation pattern. The local clustering of cell types by their epigenomic profiles in IDEAS allows the system to learn the signal distribution for a feature missing in one cell type from the available signal in locally related cell types, and then use that signal distribution when assigning likely states in the cell type with missing data. While full determination of all biochemical features in each cell type is preferred, attaining complete coverage is difficult, especially for rare cell types. Indeed, many integrative analysis projects are contending with the challenges of missing data (Ernst and Kellis 2015; Schreiber et al. 2020; The ENCODE Project Consortium et al. 2020). We suggest that the IDEAS method provides a principled approach with good utility for integrative analyses in the face of missing data.

Our collection of cCREs in mouse blood cells efficiently captures known erythroid regulatory elements and potential enhancers predicted by available EP300 occupancy data. However, this initial cCRE registry is unlikely to be complete, especially for cell lineages underrepresented in our collection. The VISION resources can be useful for analysis of new data from users, such as searching for overlaps of the cCREs with peaks from new datasets. Also, parallel efforts, such as the Immunological Genome Project (Yoshida et al. 2019), are generating complementary resources that can expand the cCRE registry. Only DNA intervals in nuclease accessible chromatin were assigned as cCREs, and thus, any regulatory elements that function in nuclease inaccessible regions will be missed. Such elements may be discovered by further studies on inaccessible regions that are bound by transcription factors. Given the absence of comprehensive reference sets of known regulatory elements, neither the completeness nor the specificity of the cCRE collections can be evaluated rigorously. Future work evaluating experimentally the impact of cCREs on gene expression will provide a more complete assessment of the quality of the registry.

Each cCRE has been annotated with its epigenetic state in each cell type and an initial estimate of the epigenomic regulatory potential (eRP) score for regulating candidate target genes. These initial eRP scores for cCREs, derived from a multivariate regression and sub-selection procedure, can explain a substantial portion of variance in gene expression, but a considerable amount of expression variance remains unexplained. Estimates for regulatory output could be improved by incorporating transcription factor binding site motifs (Weirauch et al. 2014), transcription factor occupancy (Dogan et al. 2015), and patterns in multi-species genome sequence alignments (Taylor et al. 2006). The target gene assignments can be refined by inclusion of data on chromatin interaction frequencies, e.g. by restricting cCRE-gene pairs to those within a topologically associated domain, or TAD (Oudelaar et al. 2017). The VISION project has analyzed Hi-C data in G1E-ER4 cells (Hsu et al. 2017) and HPC7 cells (Wilson et al. 2016) to provide coordinates of TADs (An et al. 2019) and compartments (Zheng and Zheng 2018), and our query interface allows users to use this information to refine choices of cCREs for specific genes.

The IDEAS segmentation results across cell types revealed some known transitions between states, such as poised enhancers in multilineage progenitor cells either shifting to active enhancers or losing their pre-activation signatures to become repressed or quiescent in more differentiated cells. However, one of the most common transitions has not been described previously (to our knowledge). Of the CTCF-bound sites in LSK that were also accessible to nuclease, a substantial proportion became much less nuclease accessible while retaining CTCF occupancy in differentiated cells. The reduction in accessibility reflects a change in the chromatin structure to a more closed state, but unexpectedly, the CTCF protein remains bound. Initial studies suggested that the CTCF-bound-but-inaccessible sites were associated with repressed, gene-poor regions while the CTCF-bound-and-accessible sites were enriched at constitutive TAD boundaries. However, further studies are needed to more fully investigate the functions of different categories of CTCF-bound sites.

We found a substantially larger number of cCREs in hematopoietic progenitor cells than in mature cells, with the notable exception of megakaryocytic cells. The reduction in numbers of cCREs coincides with the decrease in the size of the nucleus during differentiation and maturation after commitment to a single lineage (Baron and Barminko 2016) and a decrease in the number of genes being expressed (Fig. 4D). While this reduction in numbers of active genes and regulatory elements appears to occur in most lineages of blood cells, it was not observed in megakaryocytic cells, which retain aspects of the regulatory landscape and transcriptomes of multilineage progenitor cells. Similarity of MK to multilineage progenitor cells has been discerned previously from phenotypic similarities (Huang and Cantor 2009), transcriptome data (Sanjuan-Pla et al. 2013; Psaila et al. 2016), and global epigenetic profiles (Heuston et al. 2018). Recent studies have shown that MK cells can be derived from multiple stages of progenitor cells, including HSC, CMP, and MEP (Sanjuan-Pla et al. 2013; Psaila et al. 2016). It is intriguing to speculate that the similarity of MK to multilineage progenitor cells may indicate that multiple stages of progenitor cells could differentiate into MK without substantial changes to the regulatory landscape. Such a conservative process differs from other lineage commitment and maturation processes that involved substantial changes to the epigenome and reduction in numbers of genes expressed.

The systematic integration of 150 tracks of epigenetic data on mouse hematopoietic cells has produced an easily interpretable representation of the regulatory landscapes across these cell types along with predictions of and annotations of candidate regulatory elements. Similar systematic integration of epigenetic data in human blood cells is on-going, which will generate equivalent resources. Such resources should provide guidance on many important problems, such as suggesting specific hypotheses for mechanisms by which genetic variants in non-coding regions can be associated with complex traits and diseases (Ulirsch et al. 2016; Bao et al. 2019).

## Methods

### Cell populations and sources of epigenomic and transcriptomic data

Detailed information about the cell populations and cell lines analyzed is in Supplemental Material, section 1. The ChIP-seq and ATAC-seq procedures followed previously published methods (Wilson et al. 2010; Buenrostro et al. 2013; Wu et al. 2014; Heuston et al. 2018). Detailed information about the experimental methods, sources of datasets, bioinformatic pipelines, and quality assessments are in Supplemental Material, section 2 and Supplemental Tables.

### Data normalization and comparison

A novel method for normalization, called S3norm (Xiang et al. 2020), was used to produce comparable peaks signals without inflating background regions. This method is described in more detail in sections 3 and 5 of the Supplemental Material, and the pipeline is deposited at GitHub (https://github.com/guanjue/S3norm). The methods for comparing epigenetic signals across cell types are described in section 4 of the Supplemental Material.

### Integrative analysis and cCRE calls

The implementation of IDEAS (Zhang et al. 2016; Zhang and Hardison 2017) for the mouse hematopoietic cell datasets is described in Supplemental Material section 6, and the software is available from GitHub (https://github.com/guanjue/IDEAS_2018). The method for calling cCREs is in Supplemental Material section 8. The methods for comparing signals in peaks of nuclease sensitivity and in transcriptomes across cell types are in section 10 of the Supplemental Material.

### Estimating impact of cCREs on candidate target genes

The methodology for estimating the output of individual cCREs based on their epigenetic states and correlations with expression of candidate target genes is presented in section 17 of the Supplemental Material.

### Additional code

In addition to the pipelines for S3norm and IDEASs already mentioned in GitHub, code and scripts used in the analysis are in the GitHub repository https://github.com/rosshardison/VISION_mouseHem_code

## Data access

All raw and processed sequencing data generated in this study have been submitted to the NCBI Gene Expression Omnibus (GEO; https://www.ncbi.nlm.nih.gov/geo/) under accession number GSE143271 and to the NCBI BioProject database (https://www.ncbi.nlm.nih.gov/bioproject/) under accession number PRJNA599438. Data and servers for visualization also are available at the VISION Project website (http://usevision.org).

## Acknowledgments

This work was supported by the National Institute of Diabetes and Digestive and Kidney Diseases (grant number R24DK106766-01A1), the National Human Genome Research Institute (NHGRI) intramural funds, and NHGRI U54HG006998.

## Disclosure declarations

The authors have no conflicts of interest to declare.

## Notes

#### Summary of Updates

Major changes in the revised version were: (1) substantially shorter main text (2) further study of sites where CTCF is retained but chromatin accessibility is lost during cell differentiation (2) much material was moved to Supplemental Materials (3) Supplemental Materials was re-organized.

http://usevision.org

