## Supplemental Materials for "An integrative view of the regulatory and transcriptional landscapes in mouse hematopoiesis"

**Supplemental Material – Table of Contents**

| **Topic** | **Page** |
| --- | --- |
| **1. Sources of purified cell populations and descriptions of cell lines** | 3 |
| Supplemental Figure S1 |  |
| **2. Sources of epigenomic and transcriptomic data** | 4 |
| Supplemental Figure S2 |  |
| **3. Normalization of ChIP-seq and nuclease sensitivity data** | 9 |
| Supplemental Figure S3 |  |
| **4. Comprehensive comparison of epigenetic datasets** | 11 |
| Supplemental Figure S4 |  |
| **5. Effectiveness of normalizations** | 13 |
| Supplemental Figure S5 |  |
| **6. Major steps in the integrative and discriminative modeling of epigenetic states using IDEAS** | 14 |
| Supplemental Figure S6 |  |
| **7. Detailed description of epigenetic states learned by IDEAS on data from 20 hematopoietic cell types from mouse** | 18 |
| Supplemental Figure S7 |  |
| **8. Methods for calling cCREs** | 21 |
| Supplemental Figure S8 |  |
| **9. EP300 peaks that do or do not overlap with VISION cCREs** | 23 |
| Supplemental Figure S9 |  |
| **10. Comparisons of cCREs and transcriptomes across cell types** | 24 |
| Supplemental Figure S10 |  |
| **11. Expression of genes for hematopoietic regulators across differentiation** | 28 |
| Supplemental Figure S11 |  |
| **12. Transitions in epigenetic states of cCREs between cell types** | 29 |
| Supplemental Figure S12 |  |
| **13. Examination of state transitions in discrete groups of cCREs with the same pattern of appearance across cell types** | 32 |
| Supplemental Figure S13 |  |
| **14. Discriminative motif analysis for cCREs with different epigenetic state transitions** | 34 |
| Supplemental Figure S14 |  |
| **15. Associations between nuclease accessibility at CTCF-bound states and gene expression** | 36 |
| Supplemental Figure S15 |  |
| **16. Enrichment of CTCF-bound sites that retain or lose nuclease accessibility at TAD boundaries** | 37 |
| Supplemental Figure S16 |  |
| **17. Method for calculating epigenetic Regulatory Potential (eRP) scores** | 40 |
| Supplemental Figure S17 |  |
| **18. Epigenetic features of cCREs close the differentially expressed genes** | 47 |
| Supplemental Figure S18 |  |

**1. Sources of purified cell populations and descriptions of cell lines**

The data were gathered from a set of purified cell populations including LSK (Lin^-^Sca1^+^Kit^+^, which includes hematopoietic stem cells or HSC and multipotent progenitor cells or MPP), several multilineage progenitor cells (common myeloid progenitor cells or CMP, granulocyte monocyte progenitor cells or GMP, megakaryocyte erythrocyte progenitor cells or MEP, and common lymphoid progenitor cells or CLP), and committed cells of the major blood cell lineages at different stages of maturity (for erythroblasts or ERY and megakaryocytes or MK) (Supplemental Fig. S1). Most of the cell populations were from adult bone marrow or spleen, but some cell populations were from the hematopoietic organ fetal liver. We included data from three immortalized cell lines used extensively in mechanistic studies of gene regulation at distinct stages of differentiation and maturation. These were HPC7 cells, which are models for multilineage myeloid progenitor cells (Pinto do O et al. 1998; Wilson et al. 2010), G1E cells, which are a model for early erythroid committed cells blocked in maturation by a knockout of the *Gata1* gene, and G1E-ER4 cells, a rescued subline of G1E that partially matures to erythroblast-like cells in a GATA1-dependent manner upon estradiol treatment to activate a GATA1-ER hybrid protein (Weiss et al. 1997; Gregory et al. 1999). This collection of cell populations was heavily weighted toward the erythroid and myeloid lineages, but representatives of some of the major lymphoid lineages were included to provide a broad context for the resources built from our integrative modeling.

*Methods*

*Cell Isolation*

All primary hematopoietic cell populations were enriched from 5-8 week old C57BL6 male mice. LSK, CMP, MEP, GMP, CFUE, ERY, CFU-MK, and iMK populations were harvested and isolated from bone marrow (BM) as described (Heuston et al. 2018). Neutrophils (NEU) and monocytes (MON) were isolated from peripheral blood by as described (Heuston et al. 2018). Isolation of other cell populations was described in Lara-Astiaso *et al.* (2014).

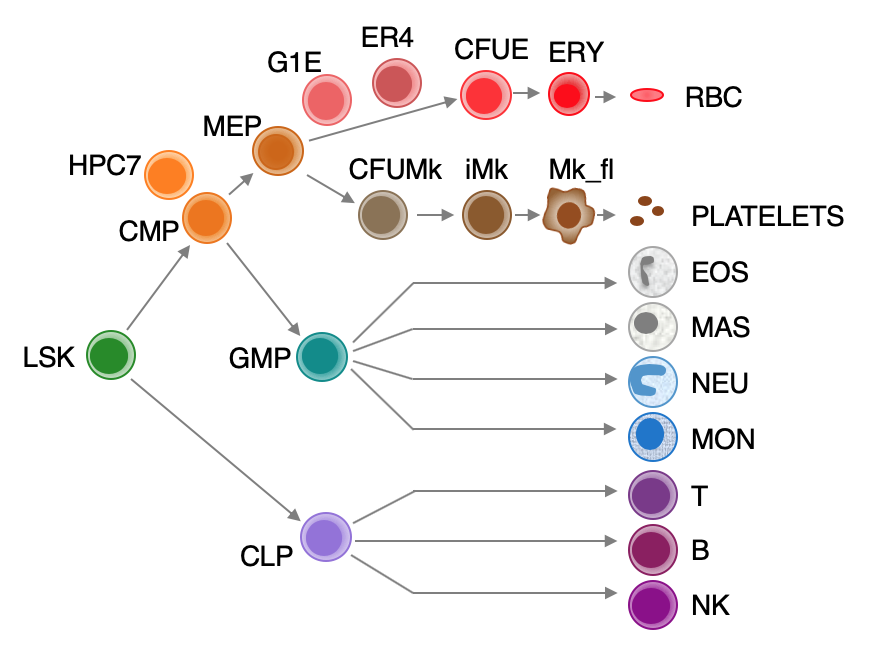

**Supplemental Figure S1.** Hematopoietic cell types. Schematic representation of the main lineage commitment steps in hematopoiesis, along with three immortalized cell lines (HPC7, G1E, G1E-ER4) and their approximate position relative to the primary cell types shown.

**2. Sources of epigenomic and transcriptomic data**

The full lists of datasets are presented in the Supplemental Tables, which is an worksheet with information about replication structure of each experiment, read counts, quality metrics, literature citations, GEO and ENCODE dataset identification numbers. An online version of this worksheet is at this link:

<https://docs.google.com/spreadsheets/d/1q7wwrTfHQlEWCq301yaF-YZk3cQMb0cRksUmgluB9kQ/edit?usp=sharing>

Results of data processing such as the signal tracks (before and after normalization), peaks for ATAC-seq and ChIP-seq data, and the estimates of transcript levels from RNA-seq (Li and Dewey 2011) are available at the VISION Project website ([http://usevision.org](http://usevision.org/)). Signal tracks, peaks, and ranges of transcript levels can be visualized at the customized genome browser at the VISION Project website.

The datasets used in this project were collected from many different sources, including individual laboratories and consortia (Supplemental Fig. S2). We started with published or previously released data from our own laboratories in both the cell lines and primary erythroid and megakaryocytic cells at various stages (Cheng *et al.* 2009; Wilson *et al.* 2010; Wu *et al.* 2011; Cheng *et al.* 2014; Pimkin *et al.* 2014; Wu *et al.* 2014; Yue *et al.* 2014; Hsiung *et al.* 2015; Jain *et al.* 2015; Stonestrom *et al.* 2015; Wilson *et al.* 2016; Heuston *et al.* 2018). However, the numbers of hematopoietic stem and progenitor cells that could be isolated by FACS on selected surface markers are small relative to those for maturing, lineage-committed cells, which presents a limitation for conventional ChIP-seq analyses. Thus, for histone modifications in these multilineage stem and progenitor cell populations, we used the ChIP-seq data obtained using the iChIP method for interrogating small numbers of cells (Lara-Astiaso et al. 2014). The iChIP data also were the primary source for epigenomic information on mouse lymphoid cells. Additional datasets obtained through the CODEX compendium (Sanchez-Castillo et al. 2015), the GEO database (Barrett et al. 2009) and the ENCODE data portal (Sloan et al. 2016; Davis et al. 2018) filled in more features for several cell types. A ChIP-seq determination on CTCF in LSK cells is a new dataset for this paper. Almost all the experiments from the VISION project and ENCODE projects were done in replicates, but experiments without replicates from any source were included if they passed quality checks (described in *Methods and Quality Assessment* below).

Several types of RNA-seq data were compiled across these cell types, including those using various strategies for RNA-seq on polyA+ RNA (Lara-Astiaso et al. 2014; Paralkar et al. 2014) and on total RNA (Heuston et al. 2018). Comparisons between the RNA-seq collections determined by different methods or using different sources of RNA (total vs polyA+) were problematic; the differences attributable to method of preparation or RNA source exceeded differences between cell populations. Thus, the set of transcriptomes determined in replicate on total RNA from twelve cell types and populations using the same procedure in the same laboratories (Heuston et al. 2018) were the primary source of expression level data used in evaluating our integration products and target gene assignments.

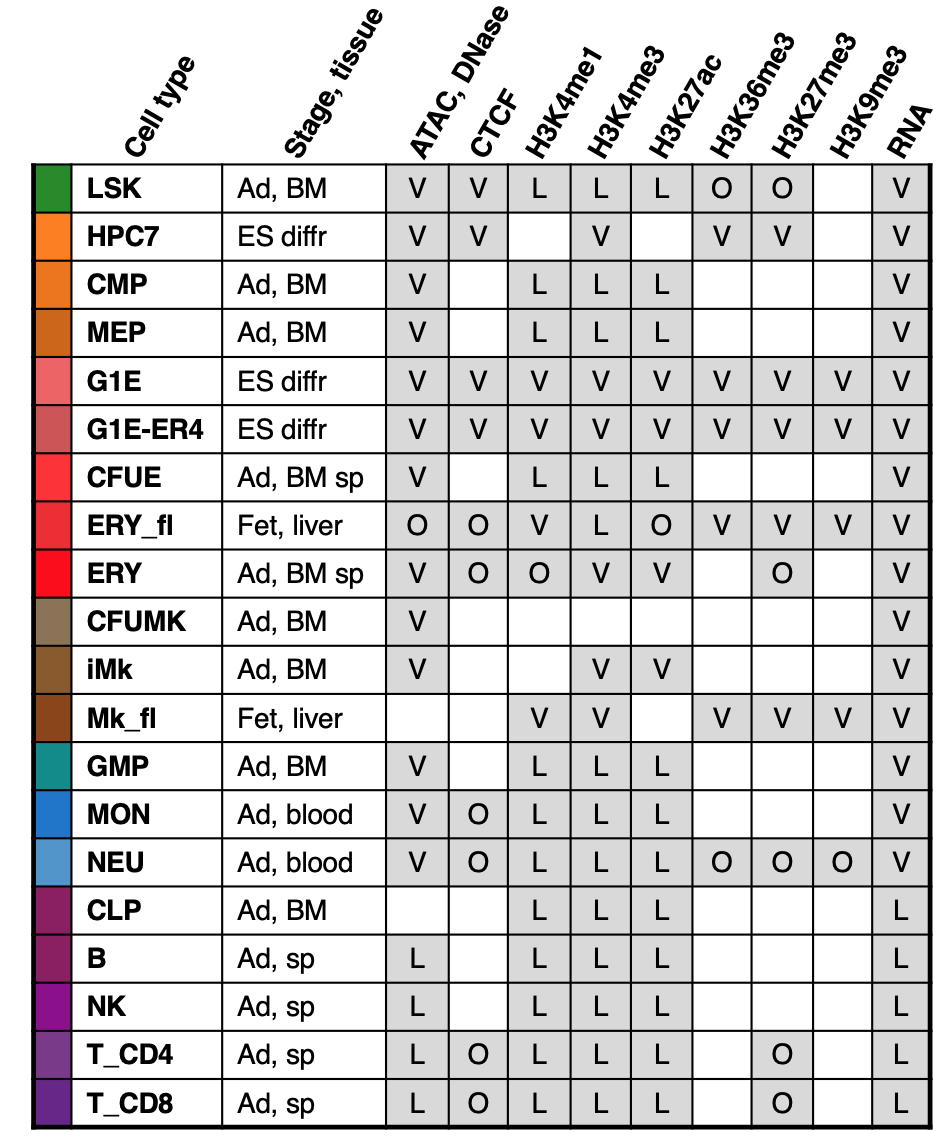

**Supplemental Figure S2.** Epigenomic and transcriptomic datasets from mouse hematopoietic cells. Across each row is presented: Cell type along with its representative color, tissue stage (Ad = adult, Fet = fetal, ES diff = Embryonic stem cell derived, differentiated) and source (BM = bone marrow, sp = spleen, liver, blood). Shaded boxes indicate the presence of the dataset, and letters denote the source (V = VISION, L = Lara-Astiaso et. al 2014, O = other). See Supplemental Table 1 for more information.

*Methods and Quality Assessment*

*ChIP-seq*

CTCF in LSK cells was performed as described (Heuston et al., 2018) using CTCF antiserum (Millipore Sigma, Cat# 07-729).

*ChIP-seq data processing*

The sequencing reads from both the new ChIP-seq experiments and many previously published ChIP-seq were processed through the VISION pipeline, mapping the reads to mouse genome assembly mm10. The VISION pipeline contains essential elements of the ENCODE mapping pipeline, but it was adjusted to allow for multiply mapping reads, which enables interrogation of duplicated genes and repetitive elements. Specifically, the VISION ChIP-seq pipeline consisted of Bowtie 0.12.8 (Langmead et al. 2009) for mapping, then filtering to remove both reads mapping to chrM or unplaced chromosomes as well as unmapped reads. The alignment was converted to bam format using Samtools 0.1.8 (Li et al. 2009). MACS 1.3.7.1 (Zhang et al. 2008) was used to generate the wiggle tracks and call peaks. The wigToBigWig program from the UCSC Genome Browser (Haeussler et al. 2019) was used to convert the wiggle file to a bigWig.

*ATAC-seq from VISION project*

ATAC-seq (Buenrostro et al. 2013) from LSK, CMP, MEP, GMP, CFUE, ERY, CFU-MK, iMK, NEU, and MON populations were generated as described previously (Heuston et al. 2018). The sequence reads were processed using a pipeline consisting of trimming the reads to 30 base pairs and then mapping using bowtie 0.12.8. The alignment was filtered to remove both reads mapping to chrM or unplaced chromosomes as well as unmapped reads. The filtered alignment was converted to bam format using Samtools 0.1.8. Bedtools 2.16.2 (Quinlan and Hall 2010) was used to convert the alignments to bed format, which were then used as input for F-Seq 1.85 (Boyle et al. 2008) to generate the wiggles. Local peaks of high frequency cleavage were determined using HOMER 1.0 (Heinz et al. 2010), as described previously (Ramirez et al. 2017).

*RNA-seq from VISION project*

RNA-seq datasets from LSK, CMP, MEP, GMP, CFUE, ERY, CFU-MK, iMK, NEU, and MON populations were generated as described (Heuston et al. 2018). The sequence reads were processed using the ENCODE3 long RNA-seq pipeline (<https://www.encodeproject.org/pipelines/ENCPL002LPE/>). In brief, reads were mapped to the mouse genome (mm10 assembly) using STAR 2.5.1b_modified (Dobin et al. 2013), followed by RSEM-1.2.28 (Li and Dewey 2011) for gene quantifications. UCSC’s bedGraphToBigWig was used to convert the bedgraph files to bigwigs for display in the browser.

*Replication and quality evaluation*

Experiments arising from laboratories in the VISION consortium and ENCODE were conducted on biological replicates (e.g. either cells isolated from different groups of mice on different days, or from the same group of mice, but collected in different aliquots from the sorter and processed separately). Experiments from Lara-Astiaso *et al.* (2014) were determined once. Information on replication structure, sequencing depth, and quality metrics are provided for each dataset in Supplemental Table 1, and the overall results are summarized in this section.

The read coverage for all experiments exceeded the recommended level (Landt et al. 2012), providing over 4.2 billion mapped reads (1,952,990,660 for RNA-seq, 1,897,113,048 for ATAC-seq, and 353,030,332 for ChIP-seq) supporting the results of the experiments. The data were high quality, as evaluated by metrics currently recommended by the ENCODE Project Consortium. All ATAC-seq datasets had a FRiP score of 0.27 (27%) or greater, and all ChIP-seq datasets had a FRiP score of 0.03 (3%) or greater, consistent with the currently accepted ENCODE standards for ATAC-seq, as described at <https://www.encodeproject.org/data-standards/>. All DNase-seq datasets had a FRiP score of 0.21 (21%) or greater.

The replicates within RNA-seq experiments were highly correlated, with Spearman correlation coefficients equal to or greater than 0.93 for almost all experiments. The exceptions were RNA-seq for CFUE and ERY, for which the replicate correlation was 0.89. This slightly lower correlation values may result from the fact that RNA isolated from these cell types exhibited consistently lower RIN scores, which may reflect the presence of older, degraded transcripts due to the nuclear condensation process during erythroid maturation.

**3. Normalization of ChIP-seq and nuclease sensitivity data**

The datasets for ChIP-seq, ATAC-seq, and DNase-seq came from heterogeneous sources with considerable differences in sequencing depth and signal-to-noise ratio. We developed a new method, called S3norm, to simultaneously adjust for both sequencing depth and signal-to-noise ratio (Xiang et al. 2020). In brief, the S3norm method converts raw signal (number of mapped reads per genomic interval) to p-values, selects reference datasets to use as standards, and normalizes via a nonlinear transform to adjust background and foreground simultaneously. This latter adjustment was effectively accomplished by rotation of a regression line through a scatter plot of signal strengths for each window in the dataset being normalized and the proxy reference, such that the mean signals for common peaks were the same between normalized and reference datasets, and the mean signals for common backgrounds were also the same for the two datasets (Xiang et al. 2020). In practice, this genome-wide normalization using S3norm adjusted peaks heights to be comparable across datasets without inflating the backgrounds (Supplemental Fig. S3). To maintain a balance between replicated data for some cell types and non-replicated data for others, replicate data were merged after conversion to p-values so that only one dataset was used for each feature in each cell type. The S3norm pipeline is available from GitHub at the link [https://github.com/guanjue/S3norm](https://nam01.safelinks.protection.outlook.com/?url=https%253A%252F%252Fgithub.com%252Fguanjue%252FS3norm&data=02%257C01%257Crch8%2540psu.edu%257C2313c7a90cd94e94f94308d71cea6eaa%257C7cf48d453ddb4389a9c1c115526eb52e%257C0%257C0%257C637009665513342292&sdata=lKN6NMt%252FA%252BuXheCIRnVQK8Vns1FDvMTXufxteUC3%252BFE%253D&reserved=0) .

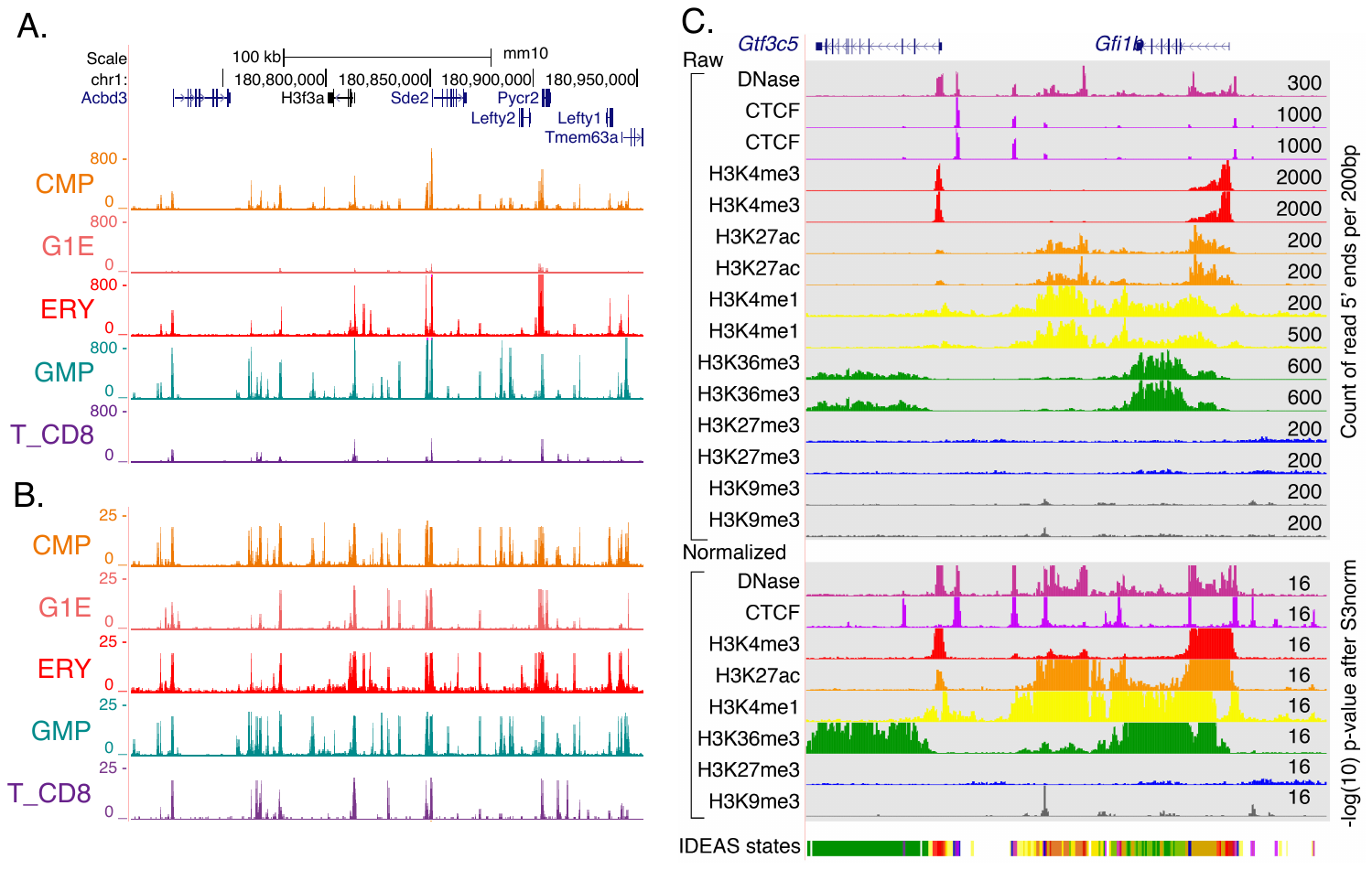

**Supplemental Figure S3.** Comparison of raw and normalized ATAC-seq/DNase-seq signal. **A.** Raw read counts for ATAC-seq/DNase-seq for representative replicates for CMP, G1E, ERY, GMP, T_CD8 at the *H3f3a* histone gene locus. **B.** Pooled, replicate ATAC-seq/DNase-seq signal for the same cell types as in (A), displayed as –log10 p-values after normalization by S3norm**. C.** Normalization across features placed all ChIP-seq datasets onto a comparable scale. As illustrated for data from fetal liver erythroblasts at the *Gfi1b* locus (70kb from chr2:28,565,001-28,635,000 in GRCm38/mm10), the raw counts of 5’ ends of reads per 200bp interval (top tracks) cover quite different scales that vary considerably among features. The maximum value is indicated at the right end of each signal track. Tracks are colored distinctively by feature (labeled at the left), and replicates are shown. After merging replicates by Fisher’s method and normalization by S3norm (Xiang et al. 2020), the signals (lower tracks of features) are displayed on the same scale, with a maximum of 16 for the negative log(10) of the p-value for deviation from a negative binomial distribution. These normalized tracks are used as input for segmentation by IDEASs, the result of which is shown for ERY_fl on the bottom track).

**4. Comprehensive comparison of epigenetic datasets**

An overview of the similarities across all the datasets showed that most clustered by epigenetic features across cell types (Supplemental Fig. S4). Results across cell types for nuclease accessibility, CTCF, the H3K9me3 heterochromatin mark, H3K27me3 Polycomb repressive mark, or the H3K36me3 transcriptional elongation mark were highly correlated. In contrast, the signature marks for promoters and enhancers, H3K4me3 and H3K4me1, respectively, formed groups interspersed with the H3K27ac modification, which is characteristic of active enhancers and promoters. These intermingled groups (e.g. H3K4me1 and H3K27ac) tended to form within related cell types, such as maturing erythroid cells or lymphoid cells, as expected for the cell type-specificity of enhancer-associated marks (Heintzman et al. 2009; Yue et al. 2014). The similarity of patterns for a particular feature across cell types suggests that examination of a single epigenetic mark may have limited power to find patterns distinctive to a cell type, whereas combinations of features appear to be more effective.

Despite our quality checks on the initially compiled experiments, four datasets were problematic. They failed to cluster with other datasets for that feature (H3K27ac for CD4 and CD8 T cells) or formed an unexpected group such as H3K27me3 with H3K36me3 for LSK (enclosed in a gray box in Supplemental Fig. S4), even after normalization. Inclusion of these datasets in the integrative modeling (described below) generated chromatin states that were highly enriched only in those cell types, unlike the other states, suggesting that they contained artifactual signals. Thus, these four datasets were excluded from the integrative and discriminative modeling.

The code and scripts used to construct heatmaps and perform other analyses are in the following GitHub repository: https://github.com/rosshardison/VISION_mouseHem_code

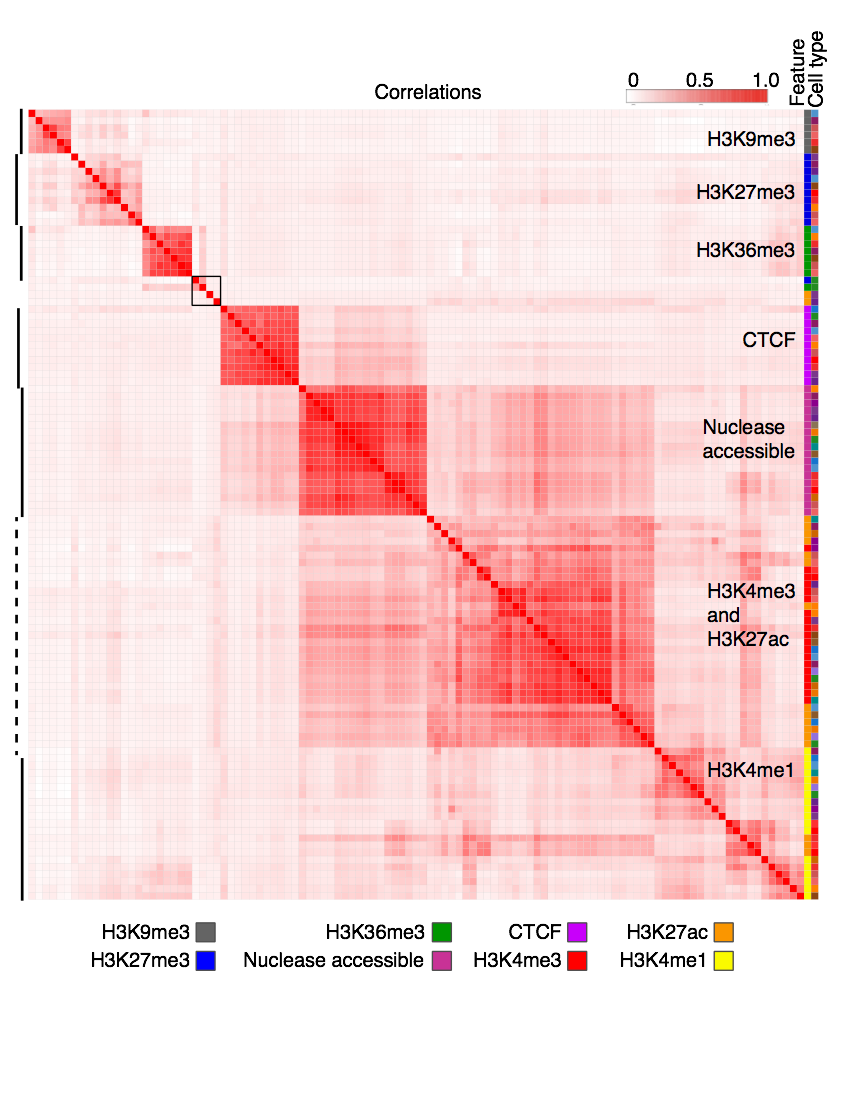

**Supplemental Figure S4.** Correlations of signal intensities across all features (S3norm normalized) and cell types. The genome-wide Pearson correlation coefficients *r* were computed for the signals in 200bp bins for each cell type-feature pair and displayed as a heatmap after hierarchical clustering (using 1-*r* as the distance measure). The features are indicated by a characteristic color (first column on right), and the cell types are indicated in the second column to the right using the same colors as in Supplemental Fig. S1**.** Four datasets failed to cluster with other datasets for that feature (gray box).

**5. Effectiveness of normalizations**

The correlation structure revealed after different types of normalization supports the effectiveness of the normalizations (Supplemental Fig. S5). The correlation matrix for the initial signal (number of mapped reads per window) showed the clustering by the feature that was emphasized with the normalized data, but it also presented a heterodisperse pattern of off-diagonal correlations and substructure between different features. The off-diagonal correlations were reduced when normalized by sequencing depth, but some clustering within cell types with similar signal-to-noise ratio was observed. Utilization of S3norm to normalize for variation in both sequencing depth and signal-to-noise ratio removed much of the off-diagonal higher correlations, which indicates that with normalization by S3norm effectively removes much of the systematic biases in the data.

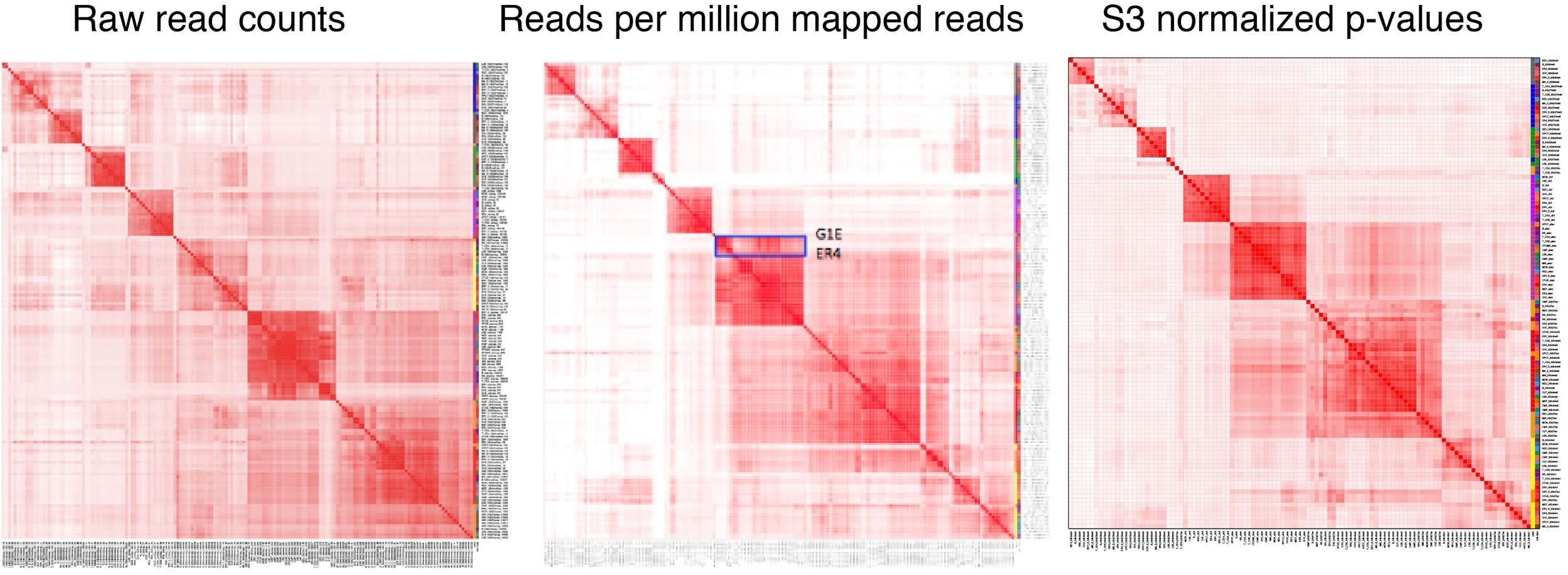

**Supplemental Figure S5.** Correlations across all features and cell types using raw data, data adjusted for sequencing depth, and data after normalization by S3norm.

**6. Major steps in the integrative and discriminative modeling of epigenetic states using IDEAS**

We employed IDEAS to analyze the normalized signals for nuclease accessibility, CTCF occupancy, and histone modifications (Supplemental Fig. S6). One key step in the IDEAS modeling is to group cell types *locally* based on the epigenomic profiles, finding regions that are similar across subgroups of cell types or that are similar across all cell types (Supplemental Fig. S6A-C). Importantly, cell types that were more similar in one locus can differ in another locus. IDEAS then learned the epigenetic states, which were defined by the signal strength, not just the presence or absence of each feature (Supplemental Fig. S6D), while retaining position-specific information (Supplemental Fig. S6E). Leveraging the information about local cell type groups and the position-specific distributions of states, genomic intervals in each cell type were assigned to an epigenetic state in an iterative process (Supplemental Fig. S6F-I).

The coloring of each state was determined automatically, generating an informative representation for each state by mixing colors from a palette of distinctive colors for each feature. The colors provide a visual representation of the contribution of each epigenetic feature to the state. The output of the IDEAS segmentation effectively paints the epigenome of each cell type in a distinctive pattern, providing a compact and concise display of function-associated states along the chromosomes of each cell type. Importantly, the segmentation provides a simplified, integrated representation of over 100 tracks of epigenomic data, enabling investigators examine the entire dataset in a concise form.

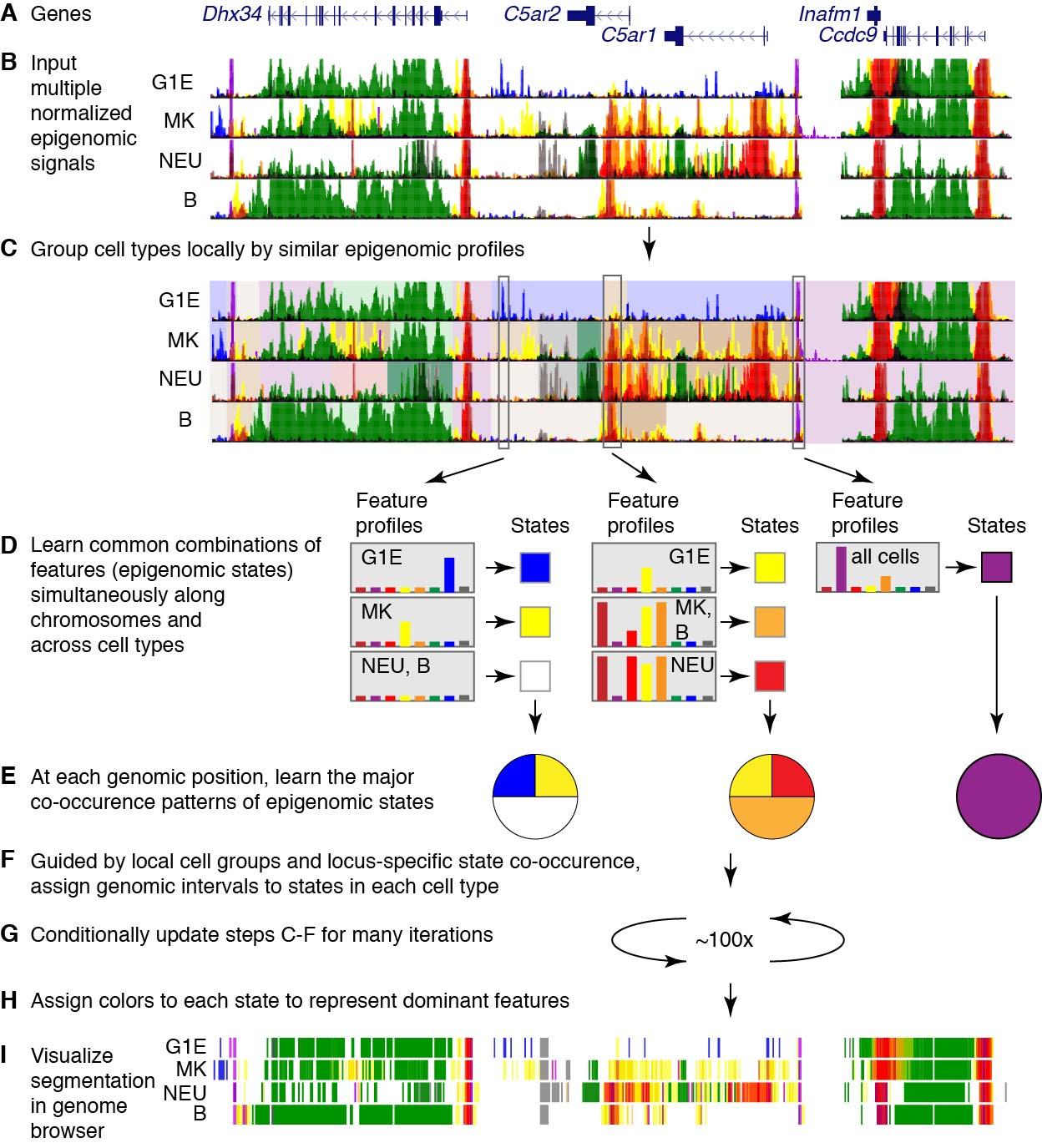

**Supplemental Figure S6.** Major steps in integrative and discriminative modeling of epigenomic signals using IDEAS. **A.** Gene models in a 100kb region centered on two complement receptor genes (position chr7:16,190,001-16,290,000 in GRCm38/mm10). **B.** In four cell types (G1E, MK, NEU, and B cells), the normalized signal for each of the eight epigenomic features was given a distinctive color (burgundy for ATAC-seq, purple for CTCF, red for H3K4me3, yellow for H3K4me1, orange for H3K27ac, green for H3K36me3, blue for H3K27me3, and gray for H3K9me3), and the eight tracks were overlaid for each cell type using the Track Collections tool of the UCSC Genome Browser (Haeussler et al. 2019). **C.** The grouping of cell types locally, based on their epigenetic profiles, is illustrated by distinctive background colors, mauve background for chromosomal segments that have similar profiles across all cell types and different colors in backgrounds for segments with differing profiles. **D.** The epigenetic feature profiles that occur most commonly are illustrated for three genomic positions as bar graphs representing the intensity of signal for each of the eight features (each with the distinctive color listed in **B**). Those combinations of quantitative signals define an epigenetic state, illustrated as a colored square. The epigenetic state at a given position can be constant or different across cell types. **E.** The frequencies of occurrence of the states at the three genomic positions are illustrated as pie diagrams; the colors in the pie diagrams represent particular states. Panels **F**, **G**, and **H** indicate steps for assigning genomic intervals to epigenetic states in each cell type and giving them informative colors. **I.** The resulting segmentation for the four cell types at this locus is shown as a track in dense mode for a genome browser.

*Methods for IDEAS segmentation*

IDEAS utilizes a Bayesian nonparametric hierarchical latent-class mixed-effect model to achieve segmentations simultaneously along chromosomes and across cell types (Zhang et al. 2016; Zhang and Hardison 2017). The computational approach has a linear time solution with respect to the number of cell types, which allows it to scale to hundreds of cell types simultaneously. For the segmentation runs described in this paper, signals in terms of numbers of mapped reads per 200 bp bin for 8 epigenetic features (histone modification and CTCF ChIP-seq, ATAC-seq or DNase-seq) were compiled from 20 cell types to produce a set of 150 tracks of data, including replicates. The replicates of the same epigenetic feature were merged (via Fishers’ method that emphasizes the replicate with the better signal-to-noise level), and signals were normalized using the S3norm procedure (Xiang et al. 2020) to generate 104 datasets. The normalized datasets were used as input for IDEAS to generate chromatin segmentation. The current version of the IDEAS method includes a preliminary, simple assessment of the most common combinations of epigenetic features to initialize the model building. We first binarized the signals of each feature by peak calling at FDR 0.05 using a negative-Binomial distribution as the null, where the parameters of the negative-Binomial distribution was estimated from the bottom 99% of the signals for the feature. From the combinations of 0s and 1s of multiple features at each position, we identified the distinct combinations that correspond to a preliminary set of epigenetic states. For *k* features, there are 2^*k* distinct combinations of 0s and 1s. We removed the rare combinations with <0.1% occurrence, and we used the remaining set of preliminary epigenetic states as the initial states for IDEAS. The removed rare states were replaced by a random sample of the common states. We also applied a relatively high threshold for inclusion of signals into the IDEAS modeling to produce a simpler, more interpretable model. Lowering the threshold generated many more states that were small variations on the states described here. Using the higher threshold did reduce the coverage of the genome by non-quiescent state assignments (from 19% to 14% on average). While higher coverage could be desired for some applications, for the current study we felt that the decreased coverage was off-set by the improved ability to interpret the model. The current software is available from GitHub, at the link <https://github.com/guanjue/IDEAS_2018>.

When dealing with datasets that range in quality, the IDEAS segmentation will sometimes “discover” states containing almost all features, including ones associated with opposite functions. Unlike the situation for most states, the DNA intervals assigned to such states also have high variation in the signal for each feature, indicating that the state may be further split to substates. We refer those states as ‘heterogeneous states’. When such heterogeneous states were returned, we revised our IDEAS pipeline as follows. (1) We identified the most common patterns of epigenetic peaks as introduced above from the cell types with all marks available, and we calculated the mean signal in each pattern as the initial parameters to train the IDEAS model. (2) After the first round of IDEAS run, we identified and removed the potential heterogeneous state and used the remaining states as priors to retrain IDEAS for a second round. The final segmentation is given by the second round of IDEAS run.

**7. Detailed description of epigenetic states learned by IDEAS on data from 20 hematopoietic cell types from mouse**

The 27 epigenetic states resulting from the IDEAS segmentation included many expected ones, as well as others that have been less frequently studied. In this section of the Supplement, we provide more analysis of the states. The IDEAS model summary (Fig. 2A, main text, repeated for clarity as Supplemental Fig. S7 in this section) shows the prevalence of each of the eight epigenetic features as a heatmap, organized by similarity among the states. Each of the major categories of states (promoter, enhancer, transcribed, etc.) actually presented multiple states, which in turn differed in the combinations of features in ways indicative of different (but often related) functions. As summarized briefly in the main text, six states showed a promoter-like signature, with high frequency of H3K4me3 (states 18, 21, 10, 15, 24, and 11); these are displayed in different shades of red, and P is the initial character in the explicit label. These six states distinguished promoter-like signatures by the presence or absence of other features with functional implications. For instance, four promoter-like states were also nuclease accessible (states 21, 10, 15, and 24), four also had the H3K27ac mark associated with active promoters (states 18, 21, 10, and 24), one (state 24) also had CTCF, and three had the H3K4me1 modification that flanks active promoters as well as marks enhancers (states 21, 24, and 19).

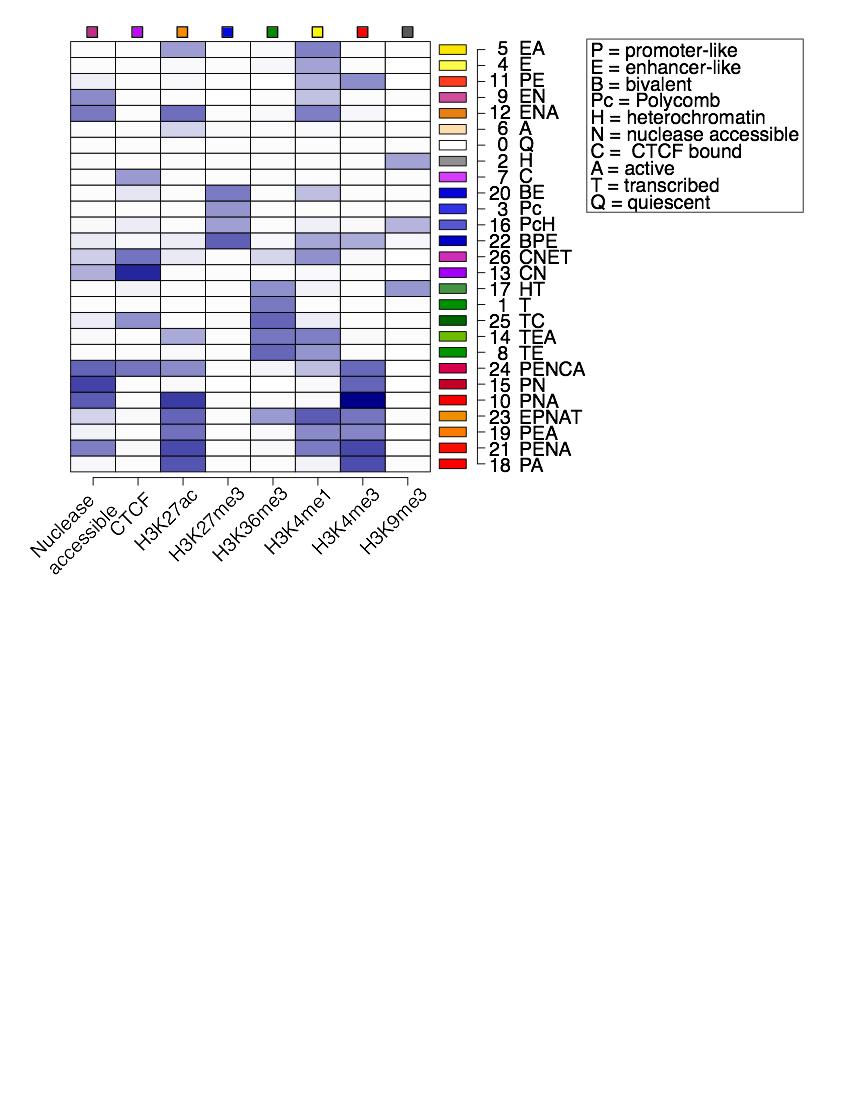

**Supplemental Figure S7**. Segmentation of the epigenomes of hematopoietic cells after integrative modeling with IDEAS. This heatmap shows the emission frequencies of each of the 27 states, with state number and function-associated labels. Each letter in the label indicates a function associated with the combination of features in each state, defined in the *box*. The indicator for transcribed is H3K36me3, active is H3K27ac, enhancer-like is H3K4me1>H3K4me3, promoter-like is H3K4me3>H3K4me1, heterochromatin is H3K9me3, and polycomb is H3K27me3.

This theme of multiple, related states capturing distinct sub-categories of major elements extended to all the major categories. In each case, the sub-category distinctions were determined by combinations of features or amount of signal for the features that suggest functional differences. Two states (19 and 23) had equivalent frequencies of the tri- and monomethylated H3K4; these were categorized as a mix of promoter-like and enhancer-like signatures, and they are colored orange. States 22 and 20 had a high frequency of the Polycomb repressive mark H3K27me3 along with methylated H3K4, and they were categorized as bivalent states (Bernstein et al. 2006). Similarly, multiple states related to CTCF occupancy (shades of purple), enhancer-like signatures (shades of yellow and orange), transcriptional elongation (shades of green), polycomb repression (shades of blue), and H3K9me3-associated heterochromatin (shades of gray) were learned in the IDEAS modeling. Several states did not fall exclusively into one of these common categories. While H3K9me3 has been associated frequently with heterochromatin, state 17 had the H3K9me3 modification together with the transcription elongation mark H3K36me3. This state is unlikely to be in repressed heterochromatin, but it is reminiscent of a previously reported association of H3K9 methylation with transcriptional elongation (Vakoc et al. 2005), a combination that was also described for KRAB-zinc finger genes (Hahn et al. 2011) and found more generally by Segway (Hoffman et al. 2012). Other states identified by IDEAS have not been previously considered in detail. One state had the expected co-occurrence of CTCF and nuclease accessibility (state 13), but an even more common state had CTCF without nuclease accessibility (state 7). While states predominated by H3K27me3 alone (state 3) or H3K9me3 alone (state 2) were common, state 16 had both repressive marks. Thus, the IDEAS segmentation learned and assigned a diverse set of states that not only included previously described epigenetic signatures but also identified some new states.

A large majority of the genome in each cell type was assigned to the quiescent, low signal state. A low-signal state covering most of the genome was observed in previous studies (Wu *et al.* 2011; Ernst and Kellis 2012; Hoffman *et al.* 2013; Yue *et al.* 2014; Roadmap Epigenomics *et al.* 2015; Zhang and Hardison 2017), but the interpretation has ranged from this representing artifacts due to high repeats and low mappability (Ernst and Kellis 2012) to a true under-representation of dynamic histone modifications, CTCF, and open chromatin in most of the genome. We favor the latter interpretation, and suggest that much of the quiescent chromatin is repressed, but in a state not subject to histone modifications that are revealed by conventional ChIP-seq. Nevertheless, the fraction of the genome in a quiescent state may be overestimated if current assays are not fully recording some modifications in chromatin (Becker et al. 2017). For example, the H3K9me3 modification in highly compacted heterochromatin may be less accessible to the antibodies during chromatin immunoprecipitation, or heterochromatin may not be sheared adequately to solubilize the compacted chromatin to produce DNA fragments that are sequenced efficiently. Even if current methods preclude the identification of some modified chromatin because of such issues, it is still the case that the DNA in the quiescent is distinctly different from that in other states.

**8. Methods for calling cCREs**

Mapped reads in the DNase-seq and ATAC-seq datasets were filtered to remove mitochondrial reads (i.e. those that map to chrM), and then peaks were called by Homer (Heinz et al. 2010). Peaks that overlapped blacklist regions were removed. For datasets that had replicates, only peaks called in both replicates were retained. The remaining peaks from all cell types were combined, and peaks overlapping by at least one nucleotide were merged. This merger caused only a modest increase the size of the peak intervals; the median sizes were 150 bp before and 263 bp after merging. The ATAC-seq or DNase-seq signal in the DNA interval corresponding to each nuclease sensitive peak in this comprehensive set was determined by aggregating the reads mapping to that interval in each cell type. Of the 215,120 merged nuclease sensitive peaks, 205,019 were not in a quiescent state (state 0) from the IDEAS segmentation. Specifically, if more than 50% of a peak interval was in the quiescent state, it was not included as a candidate regulatory element. These non-quiescent, reproducible (if replicates were available), merged ATAC-seq peaks constitute the set of **c**andidate **C**is-**R**egulatory **E**lements (cCREs).

Other cCRE datasets were obtained from the Blood Enhancer Catalog (Lara-Astiaso et al. 2014), and the SCREEN website for ENCODE cCREs (The_ENCODE_Project_Consortium et al. 2020). The SCREEN cCREs were downloaded in July of 2018, obtaining one set for all mouse cCREs and another set restricted to those with DNase-seq data for "C57BL/6 liver male embryo (14.5 days)", which are referred to as the SCREEN fetal liver cCREs.

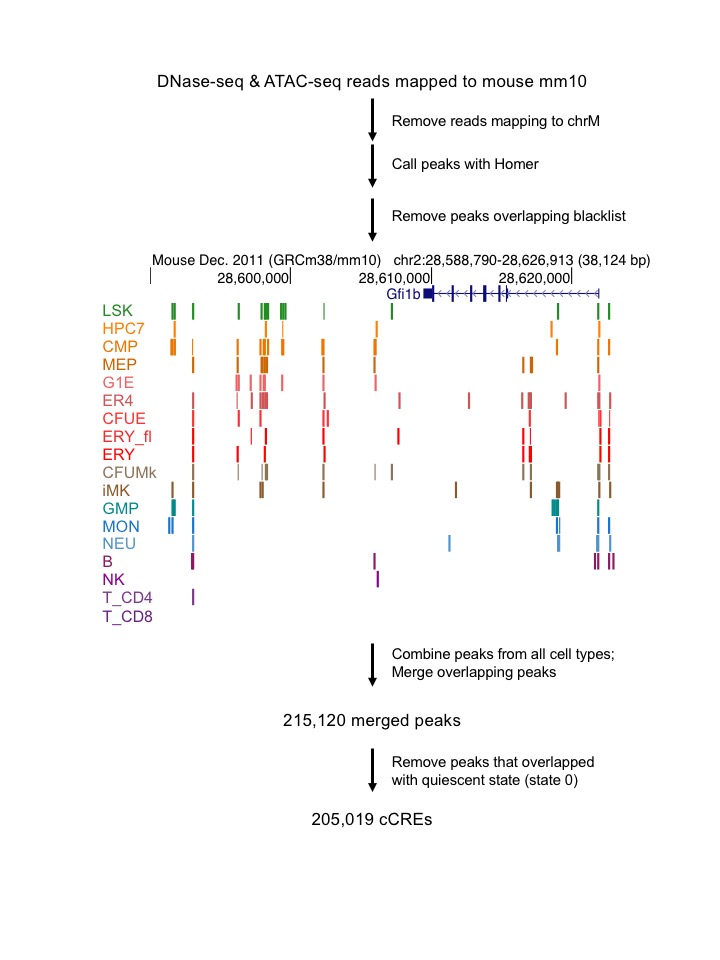

**Supplemental Figure S8**. Workflow for calling cCREs. ATAC-seq peaks (LSK, CMP, MEP, G1E, ER4, CFUE, ERY_fl, ERY, CFUMk, iMk, GMP, MON, NEU, B, NK, T_CD4, T_CD8) and DNase-seq peaks (HPC7, ERY_fl) are shown at the *Gfi1b* locus, colored by cell type. Only one replicate per cell type is shown.

**9. EP300 peaks that do or do not coincide with VISION cCREs**

While the overlap of the cCRE datasets with the collection of EP300 peaks supported the quality of those datasets, no set of cCREs captured all the EP300 peaks. This lack of full overlap raises the question of whether the EP300 peaks over-estimated the cCREs or the cCRE sets were missing regulatory elements. We examined the EP300 peaks that did or did not overlap with VISION cCREs for features that could distinguish the two groups and thus may shed some light on this issue. The signal strength of the EP300 peaks had a similar distribution for both the set that overlaps with VISION cCREs and the set that does not overlap. However, the set that overlaps VISION cCREs had a significant trend toward higher signals than did the peaks that do not overlap (Supplemental Fig. S9A). Thus, we concluded that the VISION cCREs tended to capture the stronger EP300 peaks.

Both sets of EP300 peaks were enriched for expected functional terms in various ontologies, but the set that overlapped with VISION cCREs was enriched in more terms with a greater level of significance (Supplemental Fig. S9B**)**. A subset of 10,000 EP300 peaks was randomly chosen from each set (overlapping VISION cCREs or not), and analyzed for functional term enrichment using the GREAT tool (McLean et al. 2010) Focusing on Mouse Phenotype (MGI) and MSigDB Pathways, lists of functional terms with hundreds to over a thousand terms relevant to hematopoiesis were enriched in both sets of EP300 peaks. However, the EP300 peaks that overlapped with VISION cCREs returned more terms with lower FDR Q-values when compared to the EP300 peaks not overlapping VISION. Using Mouse Phenotype as an example, peaks common to EP300 and VISION returned 1138 terms, many with extremely low Q-values, whereas the peaks only in EP300 returned 361 terms with higher, but still significant, Q-values. These distributions were significantly different (p-value<0.001 for both Student’s *t*-test and Wilcoxon test). These indicators of higher significance suggest that the EP300 peaks overlapping with VISION cCREs may be more intimately involved in hematopoietic regulation than those that do not overlap.

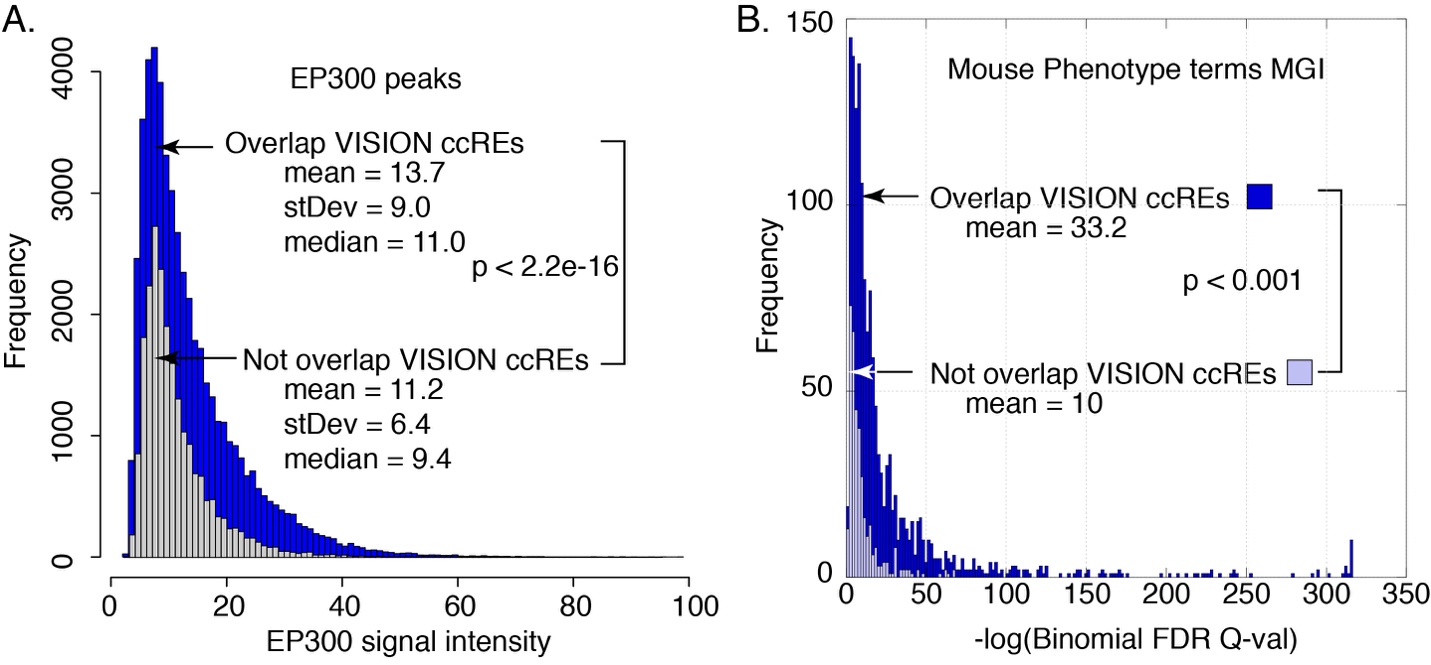
**Supplemental Figure S9.** Distributions of signal intensity (**A**) and enrichment Q-values for mouse phenotype terms (**B**) for EP300 peaks that did or did not overlap with VISION cCREs.

**10. Comparisons of cCREs and transcriptomes across cell types**

This section of the Supplemental Material presents the results of the principal component analysis (PCA) of ATAC-seq signal in cCREs and global expression levels across the hematopoietic cell populations. This section also includes the detailed Methods for these analyses.

Comparisons of the ATAC-seq signals in cCREs using the dimensional reduction approach of PCA confirmed the groupings found by hierarchical clustering, and they revealed additional insights. The first principal component (PC1) captured a substantial fraction (82%) of the variation, placing the cell types along an axis with many multilineage progenitor cells on one end and many mature cells on the other (Supplemental Fig. S10A). The values for PC1 in the nuclease accessibility analysis were strongly associated with the decrease in the number of nuclease sensitivity peaks during differentiation (Supplemental Fig. S10B, Pearson’s correlation *r*=0.92), indicating that the numbers of nuclease sensitive elements were a strong contributor to this principal component that explained a large proportion (82%) of the variation. This change in the numbers of active cCREs, with more cCREs active in the multilineage progenitor and megakaryocytic cells and fewer in other maturing lineages (Fig. 4B, main text) continued to be observed after normalizing for differences in sequencing depth, and it was robust to changes in thresholds for peak calling (data not shown). The nuclease sensitive peaks showed enrichment for histone modifications that support the accuracy of the peak calls (data not shown). The second principal component separated erythroid cells (to the left in Supplemental Fig. S10A) from other cells, and the third component tended to separate multilineage progenitor cells (toward the top) from more mature cells (toward the bottom). Thus, both the PCA and hierarchical clustering of nuclease sensitivity data in cCREs largely supported the groupings of megakaryocytic cells with progenitor cells along with separate clusters of erythroid and immune cells.

The PCA of the transcriptome data (Supplemental Fig. S10C) revealed three clusters that were consistent with the analysis of the regulatory landscape, grouping megakaryocytic cells with multilineage progenitors while keeping primary erythroid cells (CFUE and ERY) and innate immune cells (NEU and MON) in distinct groups. In contrast, MEP cells grouped with progenitor cells in the transcriptome profiles whereas they grouped with erythroid cells by nuclease sensitivity data. G1E and G1E-ER4 cell lines, which are models for GATA1-dependent erythroid differentiation, were separated from primary hematopoietic cells on PC1, and then placed between primary erythroid and myeloid cells on the PC2 and PC3 plane.

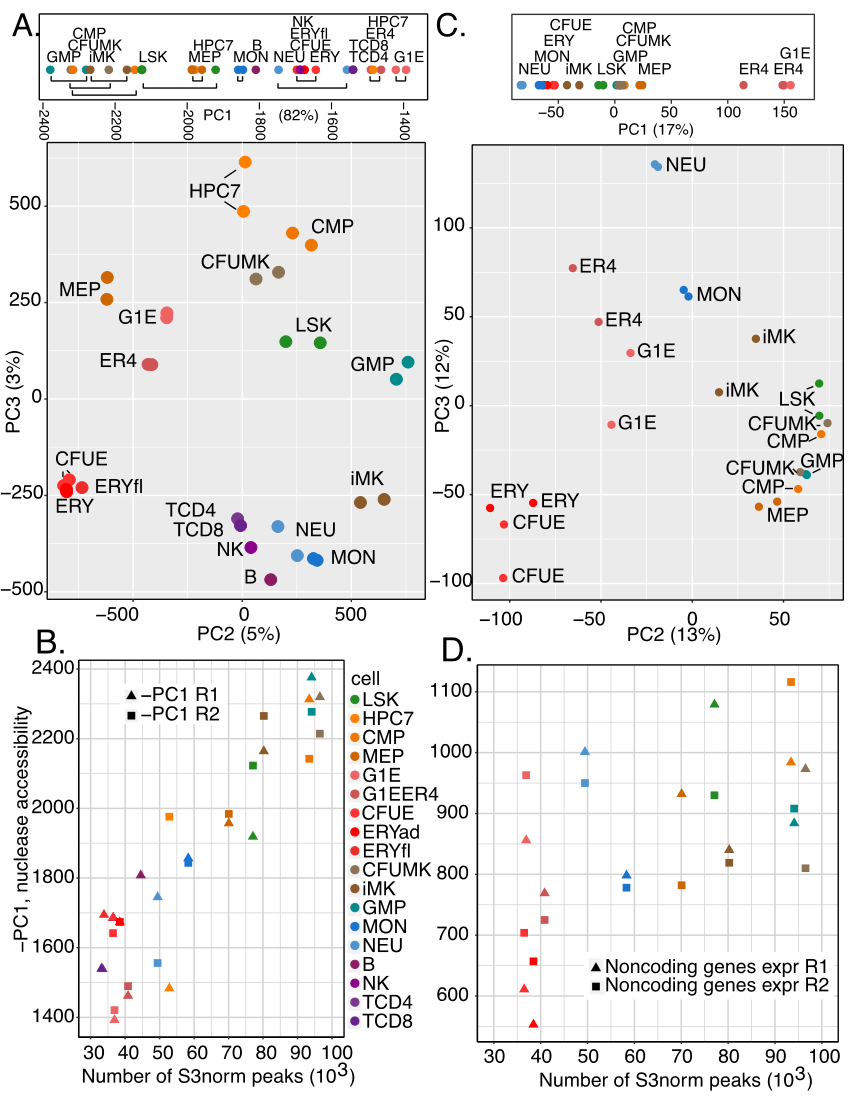

**Supplemental Figure S10.** Global comparisons of nuclease accessibility profiles and transcriptomes across mouse hematopoietic cell types: PCAs and further analysis. **A.** PCA to show groups of cell types, using ATAC-seq and DNase-seq profiles. **B.** Positive association between numbers of nuclease accessible peaks in each cell type and negative of PC1 values in PCA of nuclease accessibility. **C.** PCA to show groups of cell types, using RNA-seq. **D.** Positive association between numbers of nuclease accessible peaks in each cell type and numbers of expressed non-protein-coding genes (TPM>=1). Values for determinations in replicates are shown in panels **B** and **D**. The color code for cell types is displayed in panel **B**. R1 and R2 refer to replicates.

The numbers of noncoding genes expressed were also positively associated with the numbers of nuclease accessible peaks (Supplemental Fig. S10D, Pearson’s correlation *r*= 0.64 or 0.55 when values for G1E and G1E-ER4 cells were excluded and included, respectively, in a linear fit).

*Methods* *for* *Comparisons of epigenetic and transcriptional profiles across hematopoietic cell types*

The signal strengths of ATAC-seq and DNase-seq peaks among eighteen cell types were compared pairwise by computing the Spearman correlation coefficient (*r*) across the comprehensive set of ATAC-seq peaks for each pair of cell types, and using 1- *r* as the distance measure. Hierarchical clustering of the pairwise comparisons was performed using heatmap.2 in R.

Using the RNA-seq data on total RNA from twelve cell types (Heuston et al. 2018), we estimated transcript levels for each gene annotated by Gencode (M4) (Harrow et al. 2012), including both protein-coding and non-coding genes, using the program RSEM (Li and Dewey 2011). We compared the global transcriptomes across the cell types, again using 1 – *r* as the distance measure, and performed hierarchical clustering. The pairwise analyses reduced each comparison between cell types to a single value (r) summarizing the relationships among the ATAC-seq or RNA-seq signals. To capture the genome-wide information more completely, we also analyzed the ATAC-seq signal matrix and RNA-seq transcript levels across replicates and cell types by principal component analysis (PCA), using the tool prcomp in R.

**11. Expression of genes for hematopoietic regulators across differentiation**

The reduction in numbers of active cCREs and active genes could also lead to a reduction in the number of expressed genes encoding hematopoietic regulators. To test this hypothesis, we used the GO:1903706 category, called “regulation of hematopoiesis”, as the source of hematopoietic regulator genes, and ascertained whether they were expressed in each cell type, using three different thresholds for expression. To construct a broad set of hematopoietic regulators, we started with all the mouse genes in the GO:1903706 category, retaining 420 genes after removing redundant gene entries. Then we calculated the number of genes that have an expression level of least 1, 5, or 10 TPM in each cell type. The results showed a strong trend of decreasing numbers of expressed genes encoding hematopoietic regulators during erythroid regulation, especially at the more stringent thresholds for expression (Supplemental Fig. S11). However, no such decrease occurred in the MK, NEU, or MON lineages. A progressive decline in expressed hematopoietic regulator genes was observed for a series of mature lymphoid cells (NK > T CD8 > T CD4 > B), but the decrease was not as pronounced as the decrease in erythroid cells.

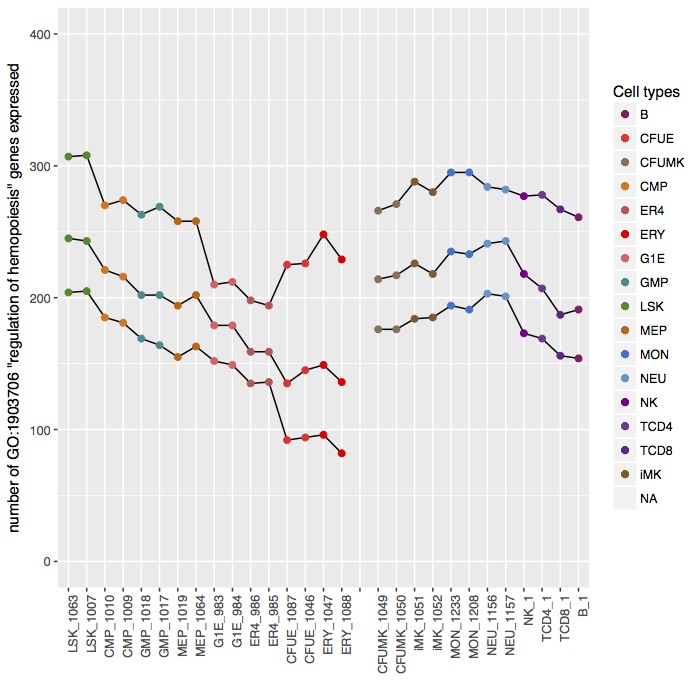

**Supplemental Figure S11**. Numbers of genes encoding hematopoietic regulators expressed in each cell type. The set of 420 mouse genes in the GO:1903706 “regulation of hematopoiesis” category was evaluated for expression using thresholds of at least 1, 5, or 10 TPM  (top, middle, and bottom lines, respectively) in each cell type. The cell types are arranged along an order of multilineage progenitor cells, differentiating G1E and erythroid cells, differentiating megakaryocytes (CFUMK and iMK), and mature cells of innate and adaptive immunity. A break in the line was introduced to emphasize the distinctive result for erythroid cells.

**12. Transitions in epigenetic states of cCREs between cell types**

While the numbers of active cCREs tended to decrease during commitment and maturation of lineages (except iMK), the reduction was particularly pronounced for cCREs in a poised enhancer mode (state 9 EN) or in a CTCF-bound nuclease accessible state (state 13 CN) (**Supplemental Fig. S12A and B**). We then determined the states into which these cCREs tended to transition by computing the enrichment for each state transition between all pairs of cell types, as illustrated for state transitions in cCREs for comparison between CMP and ERY (Supplemental Fig. S12C) and between CMP and iMK (Supplemental Fig. S12D). This examination of all state transitions in cCREs between pairs of cells revealed that the decrease in cCREs in state 9 (EN) occurred both through conversion of the cCRE to a different state and via a loss of accessibility and other epigenetic features (state 0). More specifically, cCREs in the poised enhancer state 9 (EN) in CMP tended to transition either to the active enhancer-like state 12, the polycomb-repressed state 3, or the low signal quiescent state 0 in erythroid cells (Supplemental Fig. S12C). In contrast, those CMP state 9 cCREs transitioned most frequently to state 12 (active enhancer) in iMK, with less enrichment for transitioning to the quiescent state 0 and almost no enrichment for transitioning to the repressed polycomb state (Supplemental Fig. S12D). Notably, cCREs in several different states in CMP were enriched for transitions to the polycomb state 3 in ERY (vertical blue box in Supplemental Fig. S12C). These results illustrate specific mechanisms for the recent report of more substantial changes in epigenomic landscape during differentiation of CMP to ERY than to iMK (Heuston et al. 2018).

For another major state in progenitor and megakaryocytic cells, much of the decrease in numbers of cCREs in state 13 (CTCF and nuclease accessible) occurred through a loss of accessibility while retaining occupancy by CTCF (state 7, Supplemental Fig. S12A and B). This transition from state 13 to state 7 was observed for cCREs overall during differentiation to both ERY and iMK (Supplemental Fig. S12C and D).

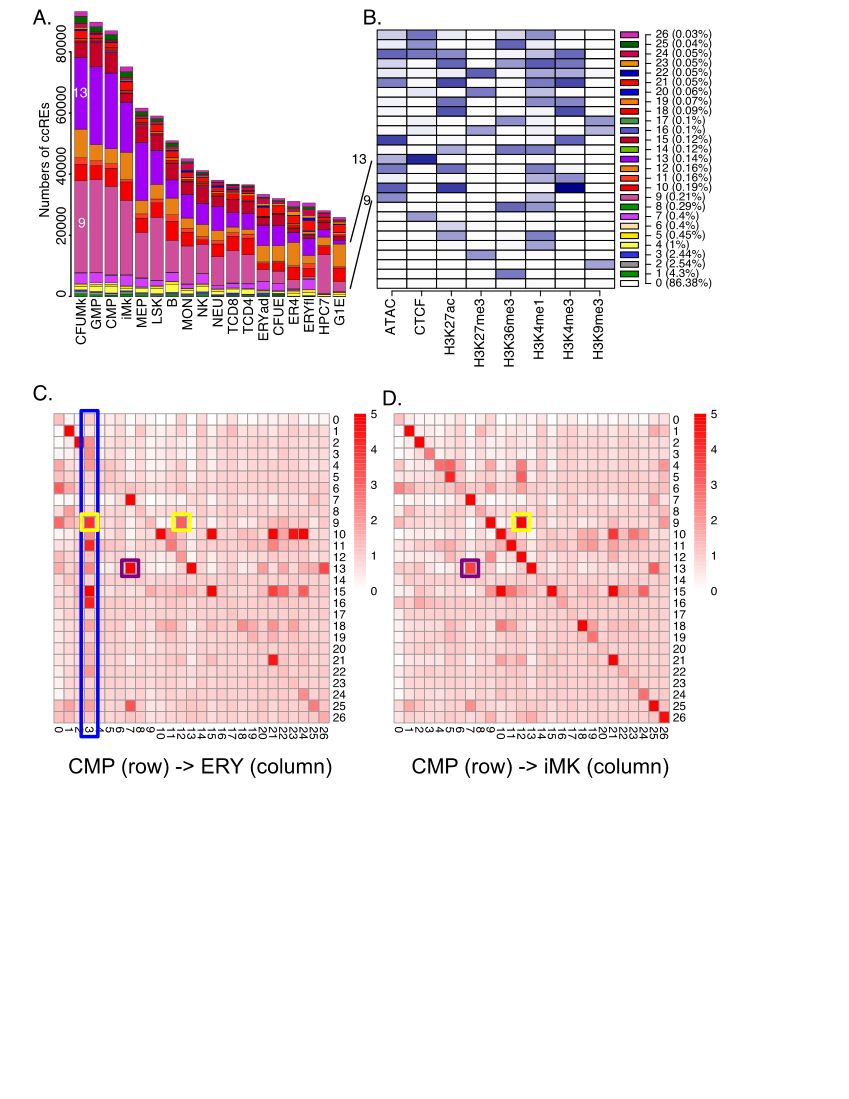

**Supplemental Figure S12.** Transitions in epigenetic states at cCREs across hematopoietic differentiation. **A.** The numbers of cCREs in each cell type are colored by their IDEAS epigenetic state, in numerical order from bottom to top of each bar. **B.** The composition of each state (heatmap in shades of blue) and the fraction of genome assigned are shown with the states in numerical order to facilitate comparison with panel **A**. States 9 and 13, which are prominent in multilineage progenitor cells, are emphasized. **C and D**. Transitions between IDEAS epigenetic states for cCREs in CMP after differentiation to ERY (C) or iMK (D). The numbers of cCREs in all state transition pairs were determined, and the enrichment was calculated as observed numbers over those expected given the numbers of cCREs in states across the whole genome in all cell types. The intensity of the red color in each cell reflects the level of enrichment. Boxes around cells emphasize transitions in states; yellow and purple for transitions from state 9 and state 13, respectively, in CMP, and blue for transitions to state 3 in ERY.

**13. Examination of state transitions in discrete groups of cCREs with the same pattern of appearance across cell types**

We employed a complementary approach to studying state transitions by examining the transitions within well-defined, discrete groups of cCREs that were clustered by their appearance pattern across cell types. The clusters were generated by an indexing approach that gives discrete categories of presence (1) or absence (0) of a nuclease accessible peak call in each cell type (Xiang et al. 2019). Thus, each cCRE had an 18-character index string, and cCREs with identical indices were placed in a group termed an index set. This clustering by discrete presence-*vs.*-absence calls across cell types gave insights into the history and progression of cCRE states.

A large fraction of cCREs in progenitor cells were in state 9 (EN, similar to a poised enhancer). Some of them transitioned to a state associated with active enhancers, exemplified by cCREs in index set 122 (Supplemental Fig. S13A). Another major trend for the state 9 cCREs was a loss of histone modifications and nuclease sensitivity to transition to state 0 (quiescent; Supplemental Fig. S12). This transition is exemplified by cCREs in index set 207, which lost nuclease accessibility in maturing cells (Supplemental Fig. S13B).

Another major state in progenitor and megakaryocytic cells is state 13 (CTCF and nuclease accessible), and the decrease in numbers of cCREs occurred through a loss of accessibility while retaining occupancy by CTCF (state 7, Supplemental Fig. S12), as illustrated by index set 165 (Supplemental Fig. S13C).

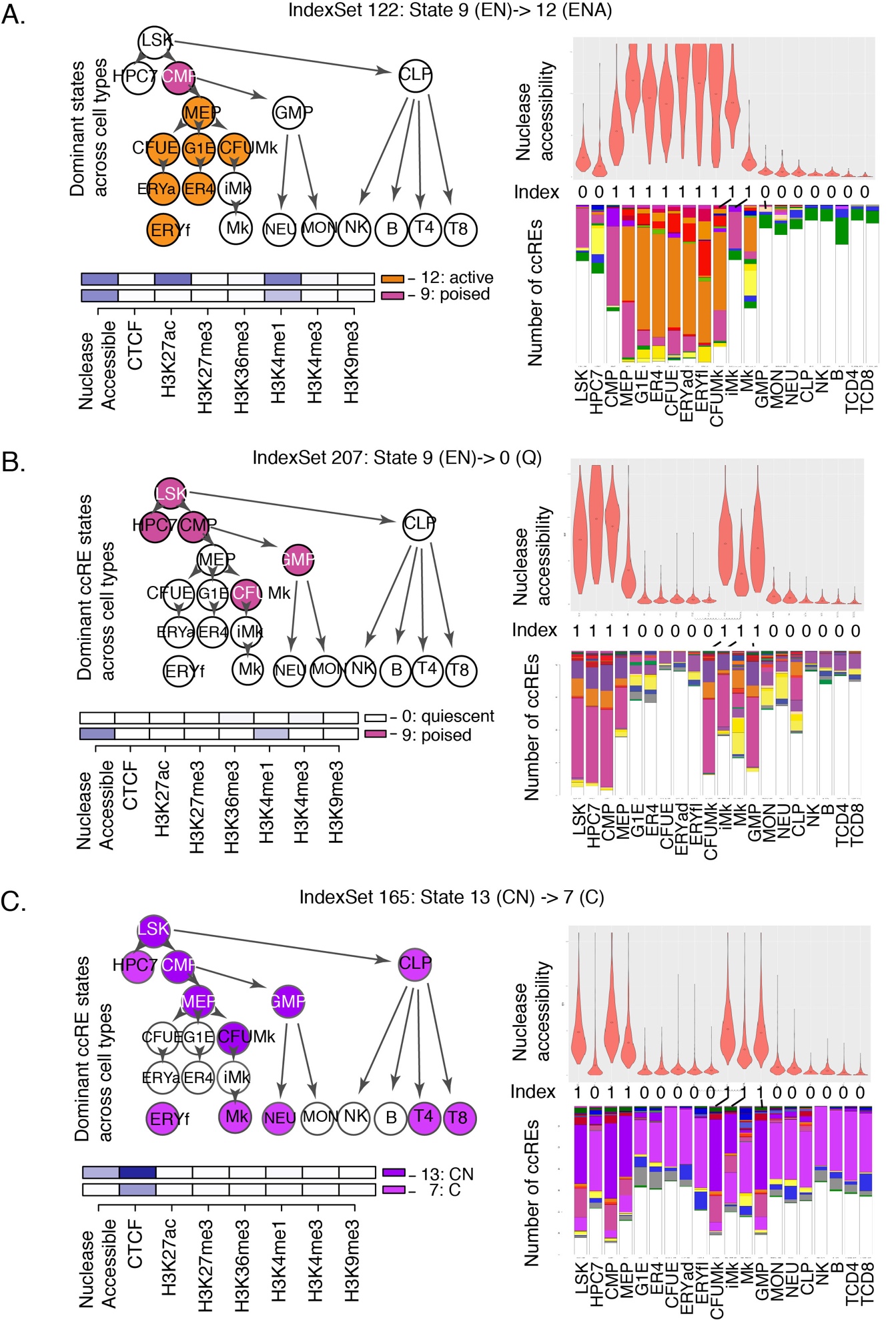

**Supplemental Figure S13**. State transitions for groups of cCREs related by their patterns of presence or absence across cell types. **A.** The state transitions and changes in nuclease accessibility across cell types are shown for the cCREs in index set 122, which undergo a transition from state 9 (poised enhancer) to state 12 (active enhancer). The index, which is a vector of presence or absence calls, is shown on the *right*, between the violin plot of nuclease accessibility and the bar plot for numbers of cCREs colored by their epigenetic states. The dominant state for this group of cCREs in each cell type is shown by the coloring of the circles in the differentiation tree on the *left*. The state compositions for the initial and final states are shown on the lower *left*. **B.** Results as in panel **A** for the cCREs in index set 207, which undergo a transition from epigenetic state 9 (EN) to state 0 (quiescent). **C.** Results as in panel **A** for the cCREs in index set 165, which are CTCF-bound sites that undergo a transition from nuclease accessible to inaccessible. These results are from the Snapshot tool (Xiang et al. 2019). Similar visualizations for all index sets can be viewed and downloaded both via our VISION website (<http://usevision.org>) or from GitHub (<https://github.com/guanjue/vision_index_set>).

**14. Discriminative motif analysis for cCREs with different epigenetic state transitions**

In CMP cell populations, we observed that the cCREs with the same poised enhancer state (state 9 EN) can change to different states such as active enhancer state (state 12 ENA) and polycomb-repressed state (state 3 Pc) in ERY cells. To explore the potential transcription factors associated with these different state transitions, we used a machine learning method called SeqUnwinder to analyzed the difference of sequence features in these categories of cCREs (Kakumanu et al. 2017). Specifically, we first extracted the cCREs in a poised enhancer state in CMP cells. The cCREs that shift to an active enhancer state in ERY cell were placed into the first group, and the others that shift to a polycomb state in ERY cell were placed into the second group. We then applied SeqUnwinder to identify the motifs that can distinguish these two groups of cCREs, using arguments “--win 400 --mink 4 --mink 5 --r 50 --x 3 --a 400 --hillsthresh 0.1 --memesearchwin 16”. The results were shown in Supplemental Fig. S14. Eight DNA binding motifs which include the GATA motif, a motif similar to the binding preference of PU.1, and additional ETS transcription factor family motifs were identified as the discriminative motifs between the two groups of cCREs. The GATA motif has a higher discriminative score for the group of cCREs that transition from poised enhancer state in CMP to active enhancer state in ERY. The PU.1 motif and the motifs of additional ETS transcription factor family members had higher discriminative scores for the group of cCREs that transition from a poised enhancer state in CMP to a polycomb state in ERY. These results suggested the different binding pattern of these transcription factors could be associated with distinct activation fates of a poised enhancer during cell differentiation.

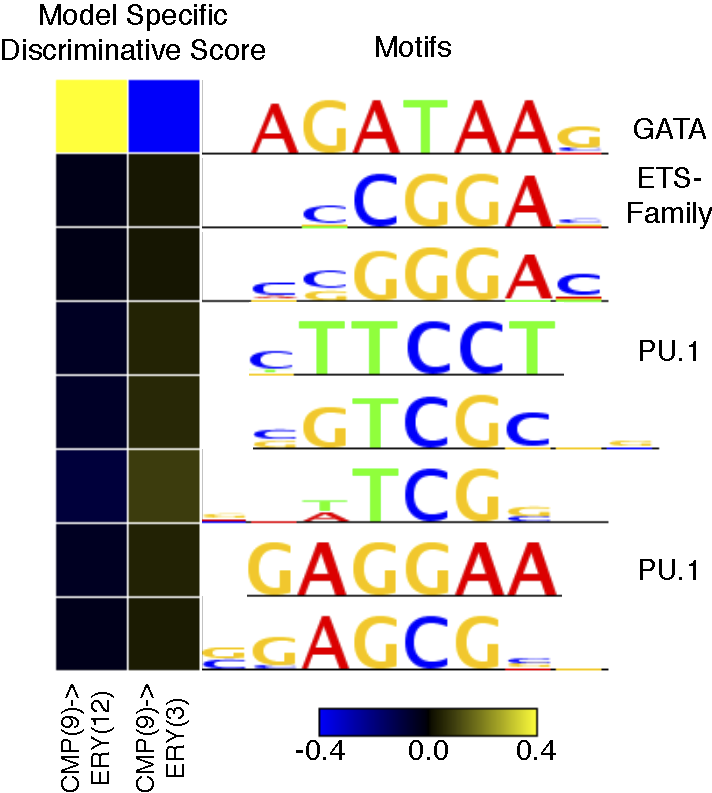

**Supplemental Figure S14.** Discriminative motif analysis of cCREs that transition from a poised enhancers state (state 9 EN) in CMP cell to an active enhancer state (state 12 ENA) or polycomb-repressed state (state 3 Pc) in ERY cell. The discriminative motifs from SeqUnwinder are shown on the right side of the figure. The heatmap on the left presents the discriminative scores of each motif.

**15. Associations between nuclease accessibility at CTCF-bound states and gene expression**

To better understand the CTCF-bound sites that lose nuclease sensitivity during differentiation, we looked for differences in expression of genes in the vicinity. We started with the cCREs with signals for both CTCF and nuclease accessibility (state 13) in LSK cells, and then compared the group that retained nuclease accessibility after differentiation to ERY (LSK 13 > ERY 13) with the group that lost nuclease accessibility (state 7; LSK 13 > ERY 7). After gathering all the genes with a transcription start site within 50kb of these cCREs, we found a consistently higher number of genes in the vicinity of the cCREs that retained nuclease sensitivity; the same result was obtained when we included genes within a longer distance (100kb) (Supplemental Fig. S15A). Focusing on the genes within 50kb of these cCREs (13,083 for LSK 13 > ERY 13 and 7,686 for LSK 13 > ERY 7), we examined the expression levels over the differentiation series LSK > CMP > MEP > CFUE > ERY. While the levels and patterns of expression varied across these genes, as expected, the genes in the vicinity of the LSK 13 > ERY 13 cCREs, i.e. the CTCF-bound sites that retained nuclease accessibility, were consistently expressed at higher levels (Supplemental Fig. S15B and C). Thus, the loss of nuclease sensitivity at CTCF-bound sites occurs in more gene-poor regions and is associated to some extent with gene repression.

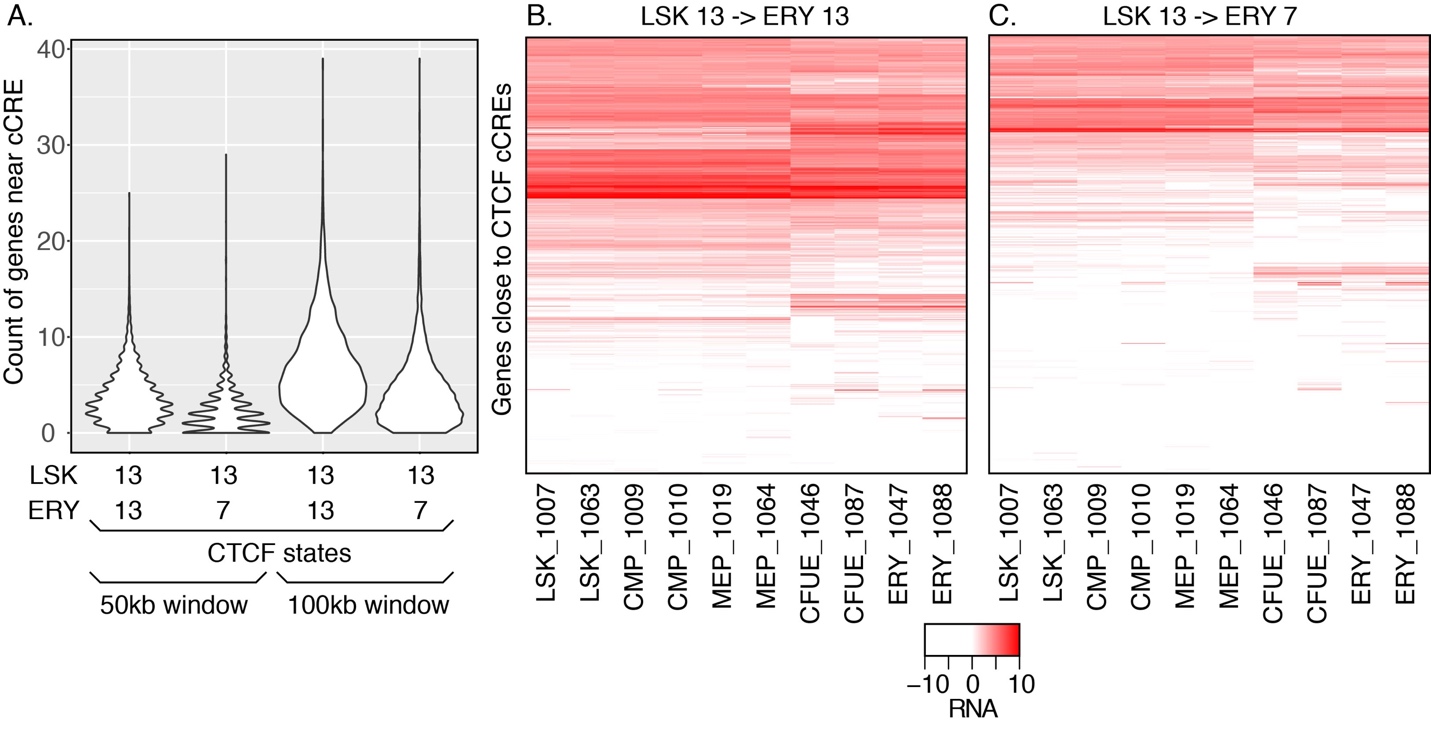

**Supplemental Figure S15**. Numbers and expression levels for genes in the vicinity of CTCF-bound sites differing in the retention of nuclease accessibility during erythroid differentiation. **A**. Numbers of genes within 50kb or 100kb on either side of each cCRE that was in state 13 (CTCF and nuclease accessible) in LSK cells, partitioned into those that retain (state 13) or lose (state 7) accessibility in ERY cells. The distribution of numbers of genes is shown as a violin plot, with the width of the violin image proportional to the number of times that each count was observed. **B. and C.** Expression levels of genes in the vicinity of CTCF-bound cCREs that retain nuclease sensitivity (**B**) or lose nuclease sensitivity (**C**) during differentiation from LSK to ERY. The expression levels (in FPKMs) of each gene (rows) are shown as a heat map across the indicated series of cell populations (with replicates), after hierarchical clustering by rows.

**16. Enrichment of CTCF-bound sites that retain or lose nuclease accessibility at TAD boundaries**

Hi-C data are available for HPC7 cells (Wilson et al. 2016), which are a cell line model for a pluripotent myeloid cell, and for G1E-ER4 cells (Hsu et al. 2017), which are a model for GATA1-dependent erythroid maturation. Thus, we could examine TAD boundaries in models for progenitors and mature erythroid cells to ascertain whether CTCF-bound sites that differed in their retention of nuclease accessibility over differentiation showed distinctive patterns of enrichment at TAD boundaries. The boundaries were called using the Optimal Nested TAD caller (An et al. 2019) at a resolution of 10kb. We then calculated the enrichment for the different categories of CTCF bound sites at TAD boundaries, with enrichment calculated by the observed number of CTCF-cCREs in a class (13_13 or 13_7) in a 10kb interval, divided by the genome-wide average for the number of CTCF-cCREs in a class in all intervals. Considering the TAD boundaries that were common to both cell lines, we found that both the CTCF-bound sites that retain nuclease accessibility (the ones that stay in state 13) and those that lose accessibility (the ones that make a transition from state 13 to state 7) were enriched at TAD boundaries. However, the ones that retain nuclease sensitivity were significantly more enriched at the TAD boundaries (Supplemental Fig. S16A). At TAD boundaries that were observed only in HPC7 cells, again both categories of CTCF-bound sites were enriched but at notably lower levels than were seen at the boundaries common to both cell types. No difference was observed in enrichment at HPC7-specific TAD boundaries between the CTCF-bound sites that retain or lose nuclease accessibility (Supplemental Fig. S16B). These results suggest that retention of nuclease accessibility at CTCF-bound sites is associated to some extent with TAD boundaries that persist through differentiation.

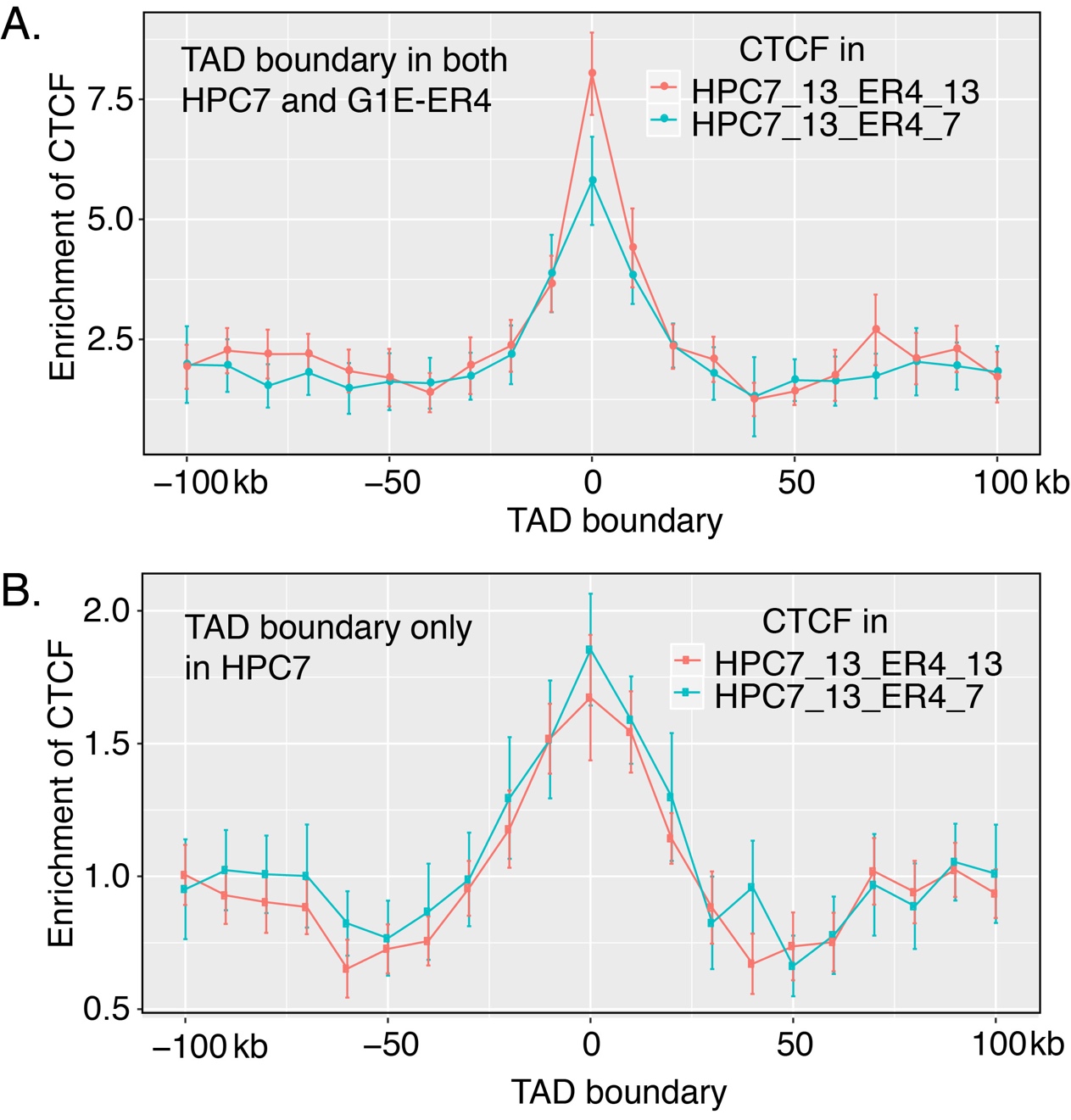

**Supplemental Figure S16**. Enrichment of two classes of CTCF-bound sites at TAD boundaries that are (**A**) preserved between HPC7 and G1E-ER4 cells, or (**B**) present only in HPC7 cells. The level of enrichment of CTCF-bound sites that were nuclease accessible in HPC7 cells and retained accessibility in G1E-ER4 cells (HPC7_13_ER4_13, red) as well as CTCF-bound sites that were nuclease accessible in HPC7 cells but lost accessibility in G1E-ER4 cells (HPC7_13_ER4_7, green) was calculated in 10kb intervals, centered on the TAD boundary and extending for 100kb on either side (x-axis). Each point is the mean enrichment at TADs across all chromosomes, and the error bar shows the 95% confidence interval.

**17. Method for calculating epigenetic Regulatory Potential (eRP) scores**

This section presents Supplemental Fig. S17, which has more detailed information on the results of the multivariate regression modeling to estimate the impact of epigenetic states and individual cCREs on expression of potential target genes. A summary of the method is given in the Results, and a detailed description is in the Methods subsection at the end of this section.

We start with our catalog of cCREs, including their epigenetic states across cell types, and expression levels of genes in 12 cell types, specifically the ones in which RNA-seq was done using the same protocol in the same laboratories (Supplemental Fig. S17A). We then use a multivariate regression approach to relate RNA-seq levels with epigenetic states of cCREs around each gene. State proportions surrounding the TSS coupled with pooled cCRE state proportions were used as predictor variables for each gene. We hypothesized that the contributions of cCREs on different types of genes are different. We thus trained the multivariate regression models using all genes or genes in four expression categories based on their average expression levels and the variance of expression level across different cell types, specifically those with (1) consistently low, (2) differentially low, (3) differentially high, and (4) consistently high expression across cell types. Within the 2Mb window considered for potential cCRE-gene pairs, there can be a large number of cCREs. However, it is unlikely that all of them contribute to the regulation of a specific gene. Thus, a sub-selection iterative routine was used to remove cCREs that contribute little to explaining expression (Supplemental Fig. S17B). The prediction accuracy in testing data after the sub-selection increased in all of the models (Supplemental Fig. S17C), which indicates that the sub-selection step effectively reduced the overfitting issue in the models. Each regression coefficient, β, from this method estimates the contribution of a specific state to gene expression (Supplemental Fig. S17D). The coefficients were computed using all genes or genes in four expression categories. The contribution of individual cCREs to expression was estimated as a weighted sum of the regression coefficients for the component states, termed an epigenetic Regulatory Potential (eRP) score for each cCRE (Supplemental Fig. S17E). Each cCRE has a specific eRP score for all potential target genes in the same category in a given cell type, but the eRP score of a cCRE differs for target genes in other categories and in other cell types.

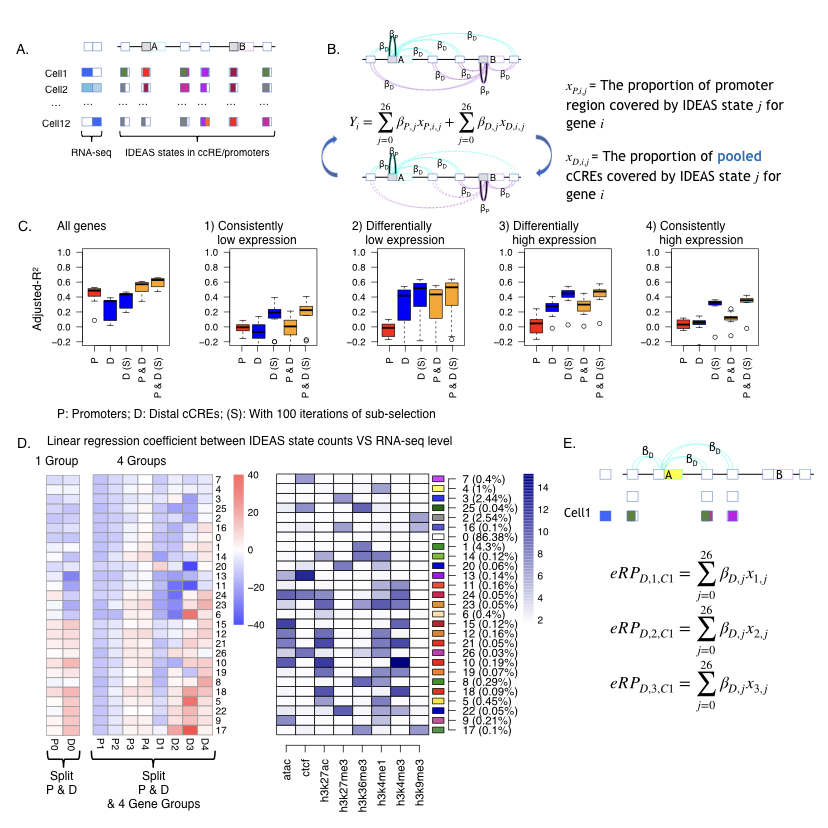

**Supplemental Figure S17.** Initial estimates of regulatory output and target gene prediction using regression models of IDEASs states in promoters and cCREs versus gene expression. **A.** Illustration of promoters and cCREs around two potential target genes, showing expression profiles of the genes across cell types (shades of blue, *left*) along with promoters and cCREs with one or more epigenetic states assigned in each cell type. **B.** Diagram illustrating the linear regression of proportion of promoters and proportion of pooled cCREs in each state against expression levels of potential target genes, learning the regression coefficients iteratively in a sub-selection strategy (indicated by dotted lines for omitted and solid lines for included cCREs in the lower diagram). **C.** Ability of eRP scores of cCREs to explain levels of expression on chr1-chr19 and chrX in the twelve cell types with and without (S) sub-selection for all genes and in the four categories of genes (1-4). A leave-one-out strategy was employed to calculate the accuracy predicting expression. The distribution of adjusted *r*^2^ values are shown as box-plots for proximal, distal, and combined cCREs. **D.** Values of the regression coefficients β for each epigenetic state for promoters and cCREs. The values of the regression coefficients for each epigenetic state are presented as a blue to red heatmap. These are aligned with a heatmap for the composition of each state, shown as shades of blue. The regressions were conducted for all genes (left two columns) or with genes in four distinct expression groups. **E.** Diagram illustrating the calculation of epigenetic Regulatory Potential (eRP) scores for cCREs as weighted sums of regression coefficients for the states covering each cCRE in each cell type.

*Methods for Mapping cCREs to Genes*

We developed a novel method to use gene expression in 12 cell types to score the cCREs for their regulatory potentials based on epigenetic states, map the cCREs to candidate genes, and further select the most likely subset of cCREs for predicting gene expression. All genes, both expressed and silent, were included so that all of the 27 IDEAS states were covered. Expecting that differences in epigenetic signals for expressed vs silent genes could affect prediction accuracy, we classified the genes into four categories: 1) consistently lowly expressed genes (mean <= -4, standard deviation or sd <= 2); 2) differentially lowly expressed (mean <= -4, sd > 2); 3) differentially highly expressed (mean > -4, sd > 2); and 4) consistently highly expressed (mean > -4, sd <= 2). For each chromosome, we used in-sample log(y+0.001) transformed tag-per-million (TPM) expression data (from all 12 cell types) to classify the genes into these four groups first, and then ran our method within each gene group and for each chromosome separately.

*1. Pre-selection of cCRE-gene pairs*

Let ***X***=(***x_0_***,...,***x***_26_) denote the proportions (between [0,1]) of each IDEAS state across eleven 200bp genomic bins, five upstream and downstream and the bin that covers the TSS of each gene. Each ***x***_i_ corresponds to the *i*th IDEAS state. Let ***Y*** denote the log(y+0.001) transformed tag-per-million (TPM) value of RNA-seq data. Note, if a gene had more than one isoform, the TSS of the largest isoform was used.

We first used the regression model ***Y***=𝛽***X***+𝜀 to obtain the coefficient 𝛽 for each state, which represents the relative impact of the *state in TSSs* on expression. After calculating the initial 𝛽s from states in TSSs, then for each gene, all genomic bins overlapping cCREs within 1Mb on either side of the TSS were assigned the TSS-derived beta value based on their state in each cell type and correlated with gene expression across cell types (excluding the left out cell type) on a bin-by-bin basis. For each cCRE, the state in each cell type of the bin with the highest correlation and overlapping the cCRE was assigned to that cCRE for that gene. All cCREs with a maximum correlation of 0.2 or higher were retained for that gene for downstream refinement. This threshold was suggested by a power curve for predicting expression, which showed increased *adjusted r^2^* values above this threshold. In addition, for any gene without a cCRE overlapping the TSS, the TSS bin was included in ***X^D^***.

2. *Refinement of cCRE-gene pairs*

Because our initial 𝛽 values are naively derived and we are using only 11 cell types for the correlation analysis, our initial application of a filter based on marginal correlation has limited power to accurately predict target genes, i.e., we do not expect all the cCREs passing that filter to be equally predictive of gene expression. We therefore developed a novel selection procedure to identify a subset of cCRE-gene pairs that are most likely capturing the regulatory relationships between cCREs and genes. Our principles are that the true cCRE-gene pairs should better predict gene expression, and that each cCRE impacts expression through their epigenetic states, thus cCREs with the same state should have the same impact on target gene expression. As such, the number of parameters in our model will only depend on the number of epigenetic states but not the number of cCREs assigned to each gene.

We assume that the impact of promoter states (within 1kb of the TSS) may be different than that of cCREs on expression. Let ***X^P^=(x^P^_0_,...,x^P^_26_)*** denote the state proportions observed at the TSS regions, and ***X^D^=(x^D^_0_,...,x^D^_26_)*** denote the state proportions observed at the cCRE regions assigned to each gene. Initially, ***X^D^*** is calculated by pooling all cCREs assigned to each gene together (based on the marginal correlation threshold), and our task is to remove some of the assigned cCREs and recalculate ***X^D^*** to maximize predictive power. Our model is in a regression form

***Y***=𝛼+𝛽^P^log_2_***(X^P^+0.001)***+𝛽*^D^*log_2_*(****X^D^+0.001)***+𝜀

where ***Y*** is the log_2_-transformed observed gene expression described above, 𝛽^P^ and 𝛽^D^ are unknown coefficients to be estimated from the model for the effects of proximal and distal elements, respectively, and 𝜀 is a Gaussian error term with mean 0. Note, however, that this is not a standard regression model: though ***X^P^*** is fixed, we will be updating ***X^D^*** by adding or removing cCREs to each gene in order to maximize predictive power. Nevertheless, given ***X^D^***, the coefficients 𝛽^P^ and 𝛽^D^ will be estimated by least squares.

After initial assignment of cCRE-gene pairs based on marginal correlation, there were on average 136 cCREs (passing the cor $\geq$ 0.2 filter) assigned to each gene, whereas there were 216 cCREs per gene without the cor $\geq$ 0.2 filter. We will inevitably over fit our model by adding or removing cCREs from ***X^D^*** if our objective is to maximize the model fitting. To examine if this is the case, we used cross-validation accuracy as a metric to evaluate our selected cCREs. Specifically, we only used data in 10 cell types to train the model but we evaluated the model performance given the current selected cCREs by the held-out cell type. At each iteration, we calculated 𝛽s, holding each of the 11 cell types out once. We then calculated mean squared errors (MSE) for each cell type using the 𝛽s for which that cell type was omitted and based on the current set of selected cCREs as a baseline MSE. Then, for each gene we cycled through all of the associated cCREs, switching their inclusion status and adding them back in or removing them from ***X^D^*** as appropriate, predicting gene expression for each cell type using the 𝛽s calculated with that cell type held out, and calculating the MSE for each cell type. All cCREs with MSE less than the baseline MSE had their inclusion status switched and added back or omitted from the current set of cCRES, as appropriate. This process was repeated for 100 iterations, stopping early if a stable set of cCREs was attained. Because multiple cCREs could be added or removed simultaneously, each change may represent a suboptimal set. We however gradually reduced the number of simultaneous cCRE changes to theoretically reach 1 per gene by the final iteration. Briefly, if more than 1000/iteration + 1 candidates were to be added or removed, only a subset, specifically 100/iteration + 1 candidates, were designated for addition or removal. Further, this reduction in the number of simultaneous changes is implemented through a random selection which weights cCREs leading to greater MSE improvement higher. Because of this, each run of the method hypothetically could produce different final selections of cCREs for a given gene.

*3. Out-sample evaluation of prediction accuracy*

To evaluate if the above method can indeed improve prediction of gene expression by sub-selecting cCRE-gene pairs, we ran the method in 11 out of 12 cell types, and then used the model to predict the gene expression in the 12^th^ cell type (not to be confused with the held-out cell type, which is one of the 11 cell types used in cross-validation). We repeated this by leaving each cell type out once, running the cCRE-gene pair selection procedure, and then finding 𝛽 coefficients by combining data from across all chromosomes and using a linear regression of the form

***Y***=𝛼+𝛽^P^***X^P^***+𝛽*^D^****X^D^***+𝜀

The resulting betas were then used to predict the expression of the 12th cell type and calculate an overall *adjusted-r^2^* of the predicted expression against the observed expression. This was done using only ***X^P^***, ***X^D^***, and both sets of predictors and with and without the cCRE selection process for each set of predictors.

4. Final 𝛽^P^ and 𝛽*^D^* calculation
To calculate the final 𝛽^P^ and 𝛽*^D^*, data was combined across all chromosomes, and two linear regressions of the form

***Y***=𝛽^P^***X^P^***+𝜀

***Y***=𝛽*^D^****X^D^***+𝜀

was used for each gene expression category or all genes together.

*5. eRP score calculation*

To calculate eRP scores for each cCRE, we combined data for all cell type-specific cCRE-gene pairs across all chromosomes and final calculated 𝛽*^D^*. eRP scores were then calculated on a gene and cell type-specific basis as the weighted sum of 𝛽*^D^* coefficients corresponding to overlapped states as follows

$$eRP_{i,n}=\overset{26}{\underset{j=0}{\sum}}\beta_{j}^{D}x_{j,n}$$

for the *i^th^* cCRE in the *n^th^* cell type across all *j* IDEAS states, where *x_j,n_* is the proportion of each state covered by that cCRE.

The code and scripts for the several steps of this method are in the following GitHub repository: https://github.com/rosshardison/VISION_mouseHem_code

**18. Epigenetic features of cCREs close the differentially expressed genes**

The multivariate linear regression to estimate the impact of each epigenetic state on target gene expression provides an opportunity to investigate important questions, such as "What are the epigenetic features of cCREs residing near differentially expressed genes in the various cell types?" To answer this question, we focused on cCREs associated with differentially highly expressed (Category 3) genes. We had calculated the regression coefficients (𝛽s) for each state with respect to expression level, as shown in Fig. 5B and Supplemental Fig. 17. We then used these results to address this issue by determining which histone modifications and other features were more frequently found in the states with a positive versus negative association. Focusing on distal cCREs, and standardizing the values for 𝛽s such that 𝛽=0 for state 0, fifteen states had positive 𝛽 coefficients for differentially highly expressed genes. After counting the number of times each epigenetic feature occurred across these positive states, we calculated the percentage of states with each feature. These positive states showed the expected features most frequently, specifically H3K27ac, H3K4me1, ATAC-seq (nuclease accessibility) and H3K4me3, while rarely emitting CTCF, H3K27me3, or H3K9me3 (**Supplemental Fig. S18**). However, some less frequently discussed states were also observed to have a positive correlation with expression, such as state 17 with both H3K36me3 and H3K9me3. As mentioned in the main text, this state with both modifications has been observed before, and the positive correlation with expression deduced from the multivariate linear regression indicates a role in gene activation. Also, a bivalent state 22 has a positive correlation with expression.

Conversely, the 10 states with negative 𝛽 coefficients had a high frequency of features expected for gene repression, such as H3K27me3 and H3K9me3 (more of the former than the latter, Supplemental Fig. S18). Several observations are worthy of further study using a more fine-grained analysis. These include (a) CTCF-containing states had a higher frequency in states with negative correlations with expression, and (b) the frequencies of ATAC-seq signal and H3K4me1 were about equivalent for states with either positive or negative correlations with expression. One might have thought that ATAC-seq and H3K4me1 would be more associated with gene activation.

The most striking difference was the pronounced higher frequency of H3K27ac in the states with positive correlation with expression (60%); it is much less frequent the states with negative correlations (20%). This confirms yet again the power of H3K27ac for predicting positively acting regulatory elements.

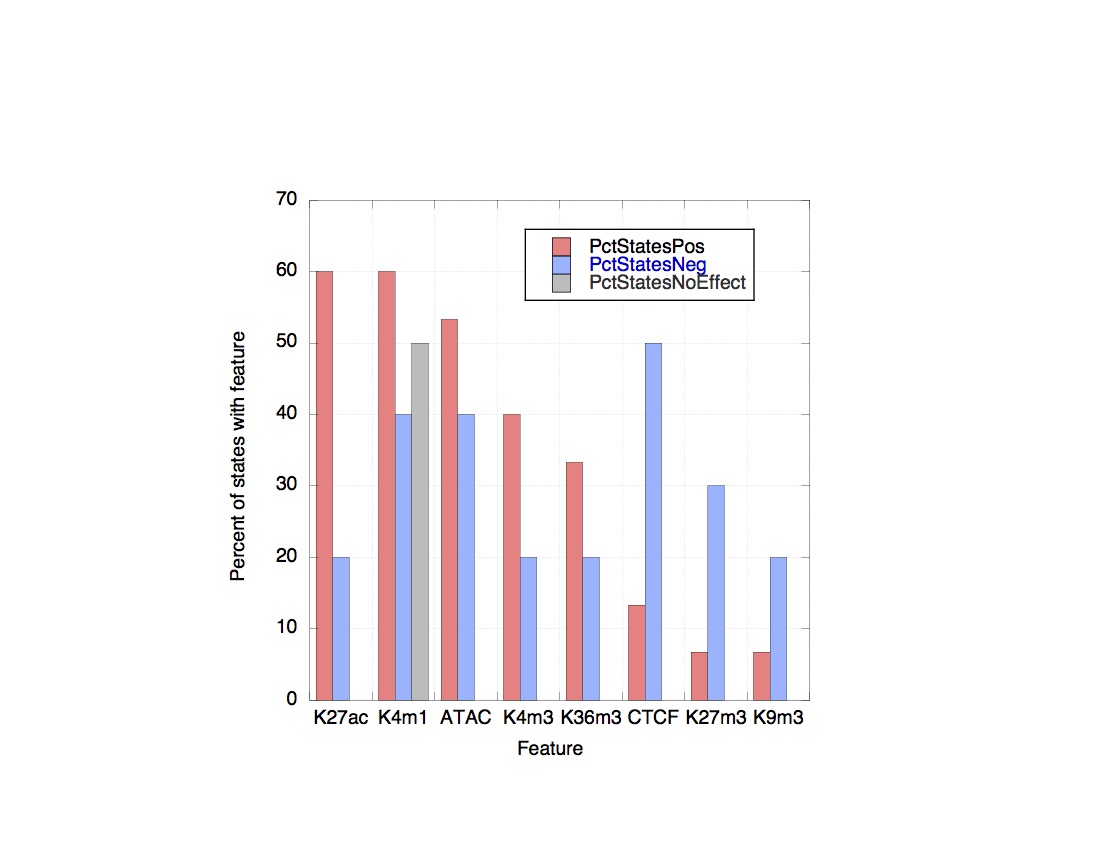

**Supplemental Figure S18.** Frequency of each epigenetic feature in states with positive or negative correlations with expression differentially highly expressed genes. The frequency of each feature is shown as the percent of states showing that feature, after categorizing the states as having a positive 𝛽 coefficient for explaining expression levels in the differentially highly expressed genes (red columns), a negative beta coefficient (blue columns), or no correlation (gray column).
